# Inflammation-associated monocytes express ACOD1 to curtail inflammatory behaviour in IBD and experimental colitis

**DOI:** 10.1101/2024.11.15.623347

**Authors:** Gareth-Rhys Jones, Broc Drury, Tobi Alegbe, Monika Krzak, Lizi M Hegarty, Tim Raine, Adam Bryne, Carl A Anderson, Gwo-Tzer Ho, Calum C Bain

**Affiliations:** Centre for Inflammation Research, Institute for Regeneration and Repair, University of Edinburgh, 4-5 Little France Drive, Edinburgh, EH16 4UU, UK; School of Infection and Immunity, University of Glasgow, Glasgow G12 8TA, UK; Wellcome Sanger Institute, Hinxton CB10 1SA, UK; Centre for Regenerative Medicine, Institute for Regeneration and Repair, University of Edinburgh, 4-5 Little France Drive, Edinburgh, EH16 4UU, UK; Conway Institute and School of Medicine, University College Dublin, Dublin, Ireland

## Abstract

Monocytes are essential for replenishing homeostatic macrophages in the intestine. However, they also accumulate in significant numbers when intestinal homeostasis is disrupted in diseases, such as inflammatory bowel disease (IBD). The molecular pathways governing monocyte behaviour across these different contexts remain poorly understood. Here, we profile the monocyte / macrophage compartment in human IBD using single-cell RNA sequencing and identify a discrete population of monocytes that accumulate in IBD, which we term inflammation associated monocytes (IAMs). These can be identified by the expression of CD319, CD274 and CCRL2, and demonstrate increased expression of IBD susceptibility genes. By performing cross-species analysis, we show an analogous population of IAMs accumulate during chemically induced colitis in mice. Using transgenic fate mapping approaches, we show these cells likely derive from a discrete precursor in the bone marrow (BM) and are locally imprinted to produce heightened IL-1β and TNF in mouse and humans. Importantly, we show that co-incident with a hyper-inflammatory phenotype, these same cells uniquely and specifically express aconitate decarboxylase (ACOD1) in response to local Toll-like receptor (TLR) and interferon (IFN) receptor signalling, to limit unrestricted cytokine production. Thus, the intestinal environment after injury instructs both inflammation and recovery specifically within a transitioning population of monocytes that are absent in health.

## Introduction

Macrophages form a dense network of cells throughout the gastrointestinal tract, playing pivotal roles in intestinal health^1^. Conversely, dysregulated macrophages are a central feature of chronic inflammatory disorders, including the inflammatory bowel diseases (IBD), Crohn’s disease (CD) and ulcerative colitis (UC). As treatments for these debilitating conditions are often ineffective, or may lose effect over time, there is an urgent need to identify cellular and molecular mechanisms that will allow better therapeutic targeting ^2–4^.

Although genetic and cellular studies have suggested that no single cell type may be responsible for triggering IBD^5–7^, a growing body of work indicates that monocytes and their macrophage progeny form a key nexus in the pathology^1,8,9^. First, monocytes and macrophages accumulate in vast numbers in active IBD, where they are potent producers of pro-inflammatory cytokines, including TNF, IL-1β, IL-6 and IL-23^10–12^, many of which are targets of current IBD therapies. Next, genetic or pharmacological blockade of monocyte recruitment is known to ameliorate inflammation in a number of experimental colitis models^13–17^. In addition, many IBD susceptibility genes are associated with monocyte and macrophage function, suggesting that genetic dysregulation of these genes might increase the risk of developing IBD^18,19^. As such, there is a great need to understand the heterogeneity of the intestinal monocyte and macrophage compartment in both health and IBD.

A number of studies have suggested that the pro-inflammatory functions of tissue macrophages in IBD are due to monocyte-like cells that are phenotypically and functionally distinct from their homeostatic counterparts^9–11,20,21^. Indeed, a persistence of these cells has been correlated with anti-TNF treatment failure^22^. However, given that monocytes have been shown to continually enter the intestinal mucosa to replenish resident homeostatic macrophages^15,23–25^, it remains unclear how the same progenitor cell can give rise to such functionally distinct progeny. Furthermore, monocyte recruitment is thought to be crucial for mucosal repair following inflammation, but the molecular pathways that control monocyte behaviour and allow a switch in their function remain poorly understood.

Here we have performed scRNA-seq analysis of monocytes / macrophages in the context of human CD and experimental colitis to identify a novel population of monocytes defined by *CXCL9/10* and CD319. Using lineage tracing approaches, we show these cells likely derive from a discrete precursor in the bone marrow (BM) and differentiate locally in the intestinal mucosa where they produce heightened IL-1β and TNF in mouse and humans.

Importantly, we show that as well as inducing a hyper-inflammatory phenotype, Toll-like receptor (TLR) and interferon (IFN) receptor signalling simultaneously induce expression of aconitate decarboxylase (ACOD1), the obligate source of itaconate, which acts as a molecular ‘handbrake’ to limit unrestricted cytokine production. This behaviour is cell-intrinsic, as lack of itaconate cannot be overcome by other local sources of endogenous itaconate. Thus, the intestinal environment after injury instructs both inflammation and recovery specifically within a transitioning population of monocytes that are absent in health.

## Results

### Human inflammation associated monocytes (IAM^hu^s) are defined by *CXCL9/10* and CD319 in active Crohn’s disease

To characterise the heterogeneity of monocytes and macrophages in human intestine, we first extracted cells of the myeloid lineage from our recent single cell RNA-seq (scRNA-seq) atlas of mucosal biopsies taken from the inflamed terminal ileum of Crohn’s disease (CD) patients and healthy controls, totalling 440,000 cells from 121 individuals^26^(**Supp. Table S1**). As described in Krzak et al.^26^, cell clusters were defined using the Leiden graph-based clustering algorithm v0.8.3^26^ and a single layer dense neural network fit to predict cluster identity from gene expression using keras v2.4.3^27^. Six cell populations were identified within the myeloid cluster, which were then annotated using specifically expressed genes from CELLEX. Five of these six myeloid populations were identified as monocytes or macrophages based on their expression of *CSF1R, C1QA/B/C, S100A8/9* or *FTL* with one mixed population of *CLEC9A/10A^+^* dendritic cells that were excluded from downstream analysis (**Figure 1a & b**). Healthy control samples were dominated by *C1QA/B*^+^ mature macrophages that could be separated further based on their expression of *CD163 and ITGAX* (**Figure 1a-c**). In contrast, a population of *S100A8/9^+^* monocytes expressing high levels of *VCAN, SOD2* and *CCR2* dominated in active CD, in keeping with recent recruitment^28,29^(**Figure 1a-c**). Notably, we observed a cluster of *CXCL9/10^+^* cells expressing intermediate levels of monocyte (*S100A8/9*) and macrophage (*ITGAX/C1QA*) features that was only apparent in inflammation. The shared monocyte / macrophage identity of this cluster suggested these cells are likely in active transition from the monocyte to macrophage state (**Figure 1c**). These *CXCL9/10^+^* cells could also be defined by the expression of *SLAMF7*, *CCRL2* and *CD274,* and expressed the highest levels of many pro-inflammatory mediators, including *IL1B,* and the *ETS2* transcription factor implicated in regulating the inflammatory behaviour of macrophages in IBD^19^ (**Figure 1d**). *CXCL9/10^+^* cells were enriched for known IBD susceptibility genes, including *TNFAIP3, NOD2, IFIH1* and *JAK2,* suggesting a role in disease pathogenesis^30^ (**Figure 1e**).

**Figure 1.**
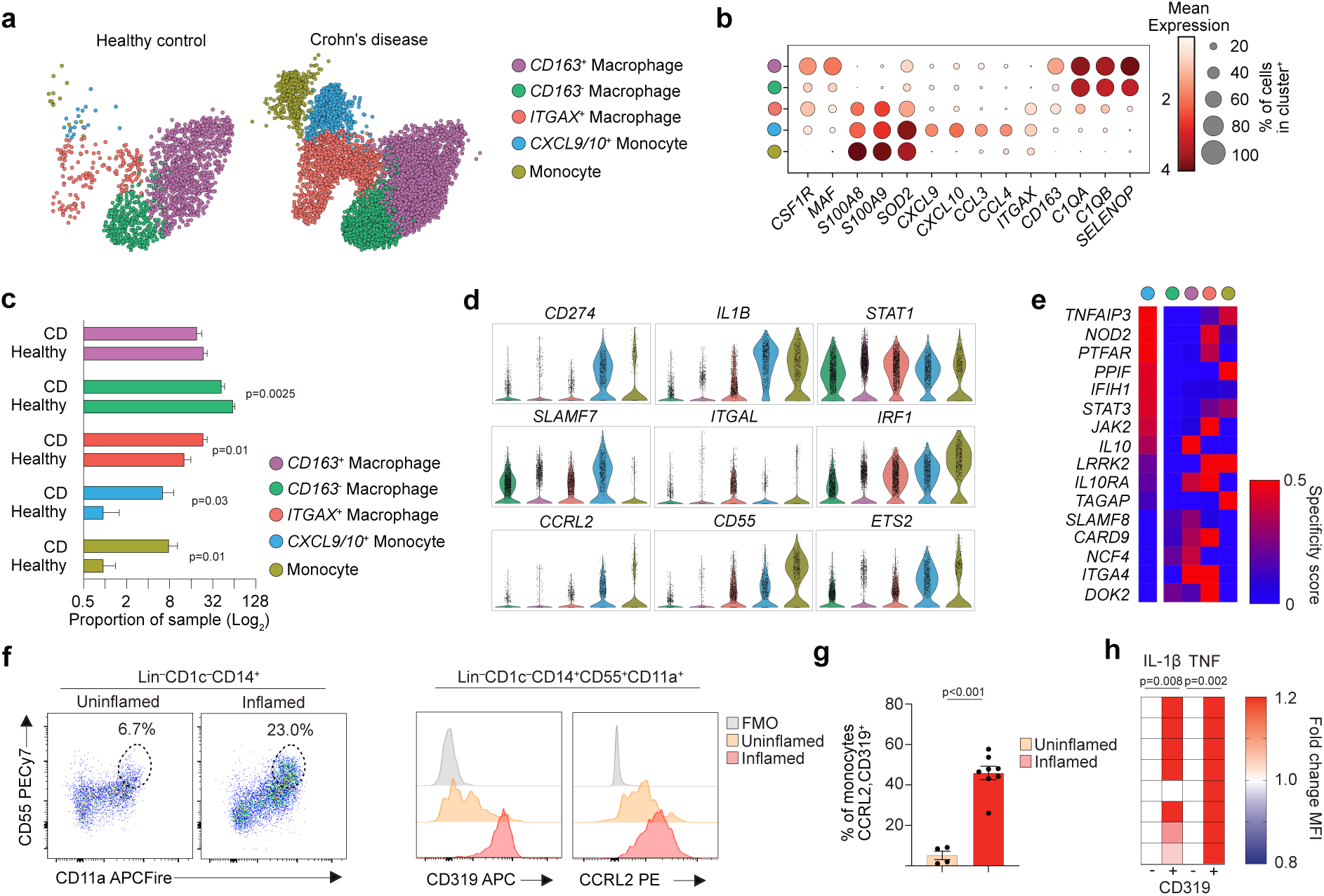
Human inflammation associated monocytes (IAM^hu^s) are defined by *CXCL9/10* and CD319 in active Crohn’s disease. **a**; UMAP dimensionality reduction of *CSF1R^+^*cells from scRNA-seq of 121 terminal biopsy samples (50 inflamed ileum from CD patients and 71 healthy controls), of which 51 patients (26 CD and 25 healthy controls) were used to generate a cellular atlas. Optimum clustering was derived from training (2/3 of cells) and test (1/3 of cells) sets, and a single layer dense neural network fit to predict cluster identity from expression using keras v2.4.3. Highly discriminative marker genes with a Bonferroni-corrected p-value <0.05 were then used to label myeloid cell types through expert knowledge. **b**; Selected canonical marker expression from a). **c**; Enumerated sequenced macrophage/monocyte cells from a), comparing CD and healthy controls. Data analysed by unpaired Student’s *t* test. **d**; Selected *CXCL9/10* cluster specifically expressed genes from a) and b). **e**; Expression of selected IBD effector genes in macrophage/monocyte cells from a) defined as; established IBD GWAS loci plus any of i) presence of a coding mutation fine-mapped down to single variant resolution, ii) detailed and convincing functional follow-up work that established the causality of the gene or iii) the protein encoded by the gene plays a major role in a pathway that is targeted by an existing IBD therapy. **f and g**; Representative flow cytometry analysis of CD55, CD11a, CD319 and CCRL2 in CD45^+^, CD3^−^, CD19^−^, HLA-DR^int^ CD14^+^ cells obtained from colonoscopic biopsies from paired macroscopically inflamed and uninflamed mucosa from 12 IBD patients. Data analysed by paired Student’s *t* test. **h**; Representative IL-1β/TNF heatmap of MFI by flow cytometry, expressed as fold change in paired inflamed-uninflamed samples from *ex vivo* cultured CD14^+^CD55^+^CD11a^+^ cells from (f).Data analysed by paired Student’s *t* test.

We next recruited an independent cohort of IBD patients and healthy controls to identify and validate transitioning *CXCL9/CXCL10*^+^ monocytic cells that we identified from transcriptional profiling (**Supp. Figure 1 & Supp. Table S2**). Targeted mucosal biopsies of inflamed, uninflamed, or healthy tissue were thus assessed for CD319, CCRL2 and CD274 surface marker expression on CD14^+^ myeloid cells by flow cytometry, using an identification strategy that incorporates both monocyte (e.g. CD55, CD11a) and macrophage (e.g. CD163, CD11c) markers^31^. CD14^+^CD55^+^CD11a^+^ intestinal monocytes were infrequent in health and inactive IBD and expressed low levels of CCRL2 / CD319 (**Figure 1g**). However, in active IBD, monocytes were significantly expanded, with the emergence of a CD319 / CCRL2^+^ subset that was not observed on CD14^+^CD55^−^CD11a^−^ macrophages (**Figure 1g & h)**. Finally, to test whether if CD319 expression identified monocytic cells with differing function, we assessed *ex vivo* IL-1β and TNF production by flow cytometry in mucosal biopsies of IBD patients and controls. Compared with CD319^−^ monocytic cells, CD319^+^ cells had higher spontaneous expression of IL-1β and TNF (**Figure 1i)**, consistent with their more inflammatory transcriptional signature.

Together, these results demonstrate that an inflammation associated monocyte population (henceforth referred to as ‘IAM’ and specifically ‘IAM^hu^’ when referring to these cells in humans), defined by *CXCL9/CXCL10^+^* and CD319, CD274 and CCRL2, expands in active IBD, and possesses superior pro-inflammatory potential compared with CD319^−^ monocytes.

### Murine inflammation associated monocytes (IAM^mo^s) are defined by *Cxcl9/10* and CD319 in acute experimental colitis

To explore the immunobiology of *CXCL9 / CXCL10^+^* monocytes further, we determined if a similar subset could be identified in murine experimental colitis. Wild type C57BL/6J mice were fed 2% dextran sodium sulphate (DSS) for six consecutive days before purified colonic CD11b^+^ / CD11c^+^ cells were analysed by scRNA-seq using the 10X genomics platform (**Figure 2a & Supp. Figure 2a & b**). We identified five clusters of *Csf1r^+^* cells from ∼16,000 high quality cells based on their expression of *Ly6c2, H2-Aa, Itgax* and *Cd163* (**Figure 2b & Supp. Figure 2c**). Whilst there was a striking accumulation of *Ccr2^+^Ly6c2^+^* cells after DSS treatment, consistent with our own and other previous work^10,15,32^, hierarchical clustering now identified heterogeneity amongst this population (**Figure 2b and c**). Strikingly, a *Cxcl9/10^+^*monocyte subset was the most expanded colon monocyte-macrophage population after DSS treatment, reminiscent of those found in human IBD (**Figure 2c and d**). We then assessed these colitis-associated *Cxcl9/10^+^*cells for the markers we identified in human IBD and found that they, like their human counterparts, had high expression of *Cd274* and *Ccrl2,* whereas expression of *Cd319* did not seem to follow the same pattern at mRNA level (**Figure 2f**). We subsequently identified and validated this *Cxcl9/10^+^* subset by flow cytometry. Akin to the human analysis, CD64^+^Ly6C^+^ colonic monocytes were relatively absent in health and expressed low levels of CD319, CD274 and CCRL2. However, during acute colitis, CD64^+^Ly6C^+^ monocytes were significantly expanded, with the emergence of a CD319, CD274, CCRL2^+^ subset (**Figure 2f)**. To assess the relative kinetics of CD319^+/–^ monocyte abundance, we then utilised a recovery model of colitis in which DSS was fed for four days before mice were returned to normal drinking water. Here, we found that the number of CD319^+^ cells peaked on day four after DSS withdrawal, parallelling the peak in histologically defined inflammation (**Figure 2g** and data not shown). Thereafter, the number of CD319-expressing monocytes diminished during resolution and repair, reaching steady state levels by day 14 after DSS withdrawal (**Figure 2g**). Furthermore, and in keeping with the human observations, CD319^+^ monocytes from mice with acute colitis demonstrated a markedly superior TNF- and IL-1β-producing capability compared with their CD319^−^ counterparts (**Figure 2h**).

**Figure 2.**
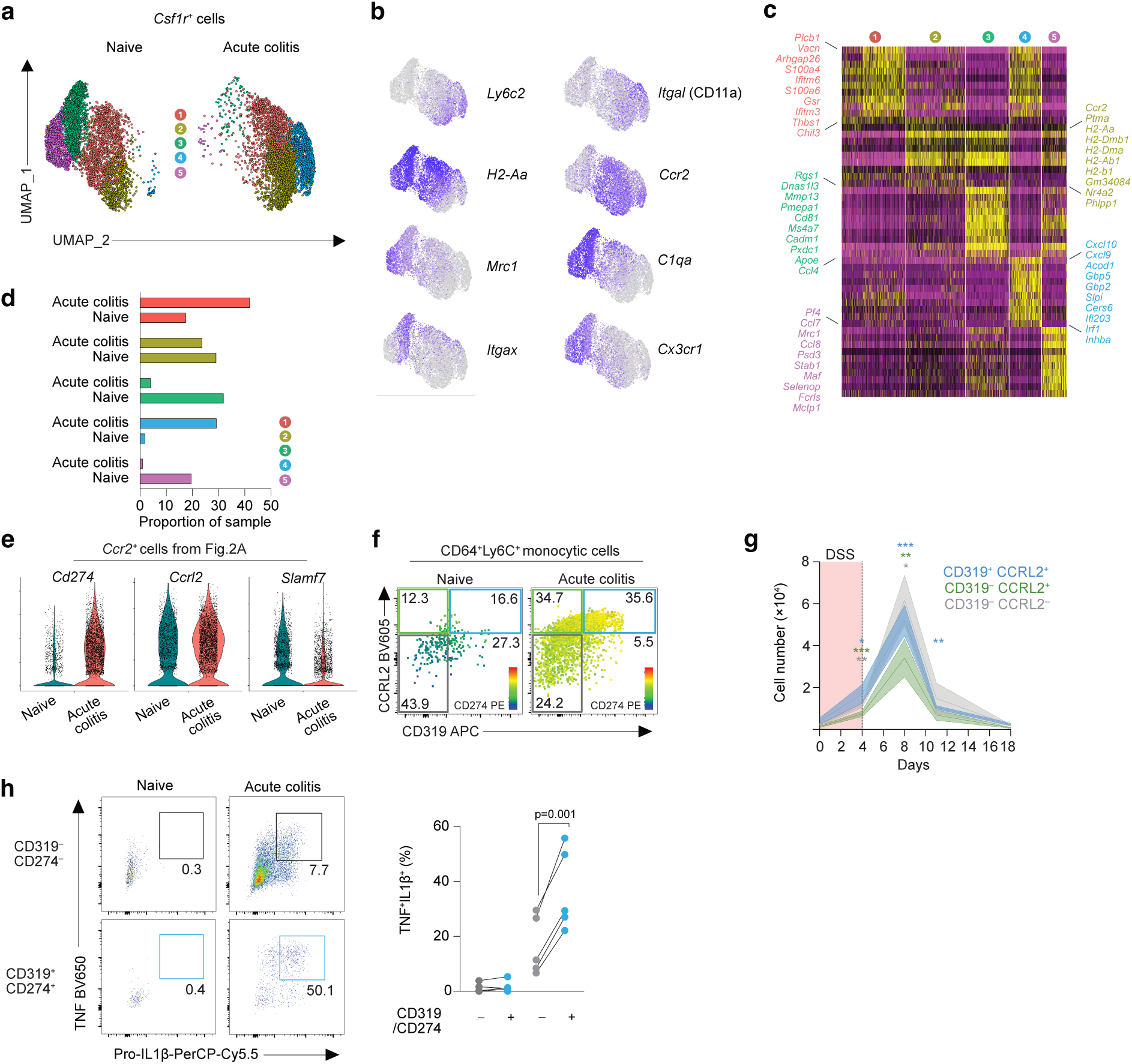
Murine inflammation associated monocytes (IAM^mo^s) are defined by *Cxcl9/10* and CD319 in acute experimental colitis. **a.** UMAP dimensionality reduction analysis of *Csf1r^+^* cells derived from scRNA-seq of CD11b^+^ and CD11c^+^ cells purified by FACS from colons of unmanipulated (naïve) C57BL/6J (WT) mice or WT mice fed 2% DSS for 6 days (acute colitis). Gating and upstream analysis of scRNA-seq shown in **Supplementary Figure 2**. **b.** Feature plots of selected cluster defining genes derived by Wilcoxon rank-sum test from data in (**a**). **c.** Enumerated sequenced macrophage/monocyte cells from (**a**). **d.** Heatmap of significant (adj *P*<0.05) top10 genes weighted by fold change, expressed in at least 25% of cells from (**a**). **e.** Gene expression of selected IAM^hu^ markers derived from Figure 1c, in *Ccr2^+^* cells from (**a**) **f.** Representative flow cytometry analysis of CD319, CD274 and CCRL2 expression in CD45^+^, CD3^−^, CD19^−^, CD64^+^, Ly6C^+^ cells obtained from colons of unmanipulated (naïve) WT mice or WT mice fed 2% DSS for 6 days (acute colitis). **g.** Absolute number of CD319 and CCRL2-defined Ly6C^+^ colon monocytes in naïve WT mice (d0) and across a recovery model of colitis induced by feeding 2% DSS for 4 days followed by 14 days on normal drinking water. Data represent n= 25, 5 per time point, from 1 time course experiment. Data analysed by paired Student’s *t* test versus control (day0). **h.** Representative flow cytometry analysis of pro-IL-1β and TNF levels (% of parent CD64^+^Ly6C^+^ monocytes after 3hours of *ex* vivo culture with 1μl/ml Monensin) split by CD319 and CD274 in naïve or 2% DSS treated WT mice for 6days (left) and enumerated (right). Symbols represent individual mice. Data are from five mice per group and are from one of two independent experiments performed. Data analysed by paired Student’s *t* test.

Thus, as in humans, a population of IAMs (denoted IAM^mo^s in mice) defined by *Cxcl9/10* and CD319/CD274 expands in acute experimental colitis and possesses superior pro-inflammatory potential compared with CD319^−^ monocytes.

### IAM^mo^s are locally imprinted and derive from a discrete precursor

To determine if the transcriptional identity of IAM^mo^s was induced locally or if intestinal inflammation led to changes in circulating monocytes prior to their arrival in the intestine, we generated a paired scRNA-seq dataset of *Csf1r*^+^ monocyte-macrophages from the blood and colon of naive (untreated) and acute DSS treated C57BL/6 mice. Using these integrated data, we applied an IAM^mo^ and IAM^hu^ signature derived from a module score of the top 20 specifically expressed genes from our human and mouse tissues (**Figure 3a & Supp. Figure S3a & b**). This showed that both the IAM^mo^ and IAM^hu^ signature was almost exclusively expressed in *Csf1r^+^* cells from the colons of mice with acute colitis and was essentially absent in cells from all other sources (naïve colon and blood), suggesting this phenotype was imprinted in monocytes only after arrival into the inflamed colon (**Figure 3b)**. Consistent with this, the expression of CD319, CD274 and CCRL2 was higher in Ly6C^+^MHCII^+^ monocytes, which represent transitioning monocytes, than in newly extravasated Ly6C^+^MHCII^−^ monocytes^10^ (**Figure 3c**).

**Figure 3.**
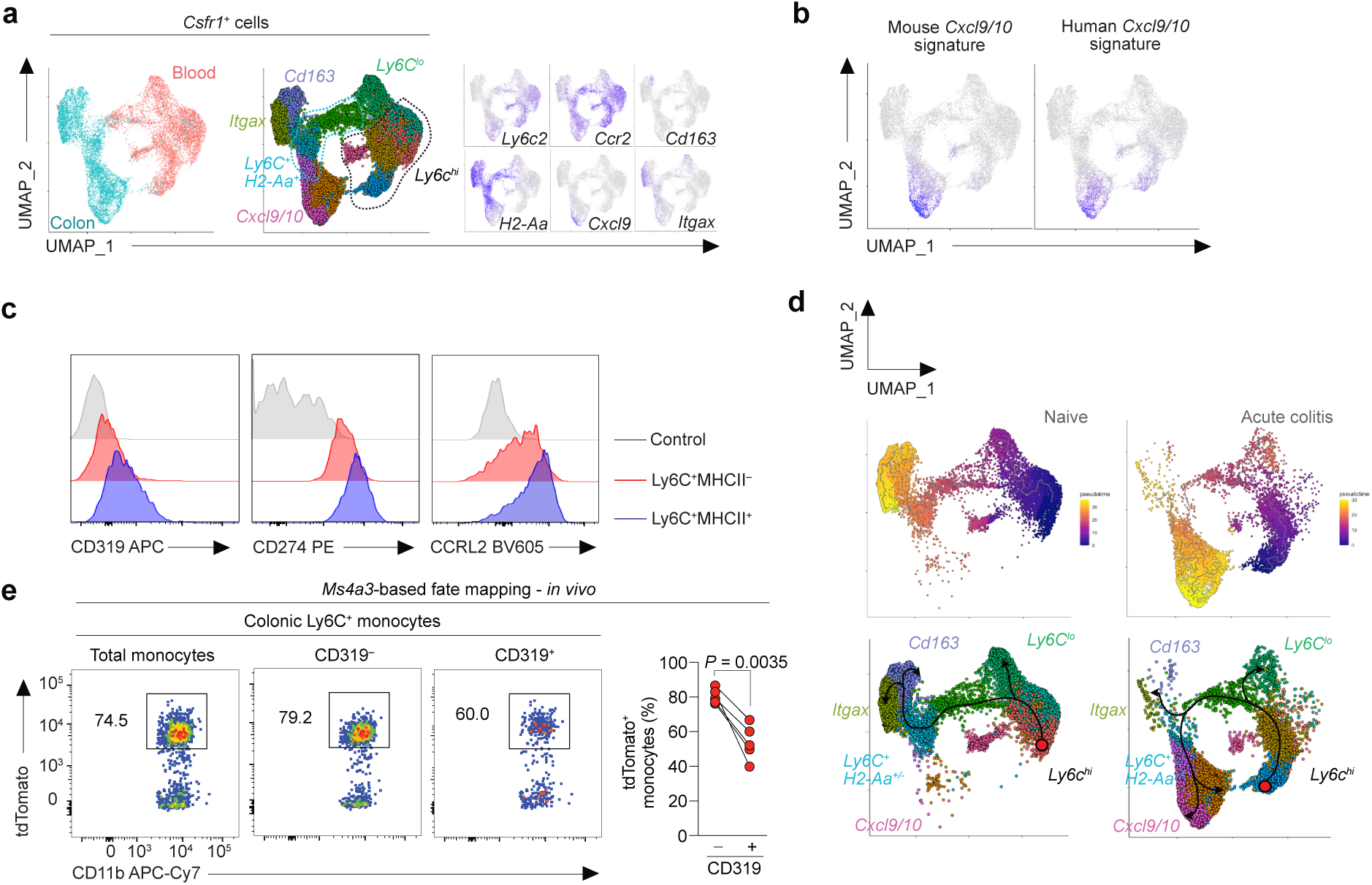
IAM^mo^s represent a discrete subset of monocytes. **a.** UMAP dimensionality reduction analysis of paired colonic and blood *Csf1r^+^* cells isolated from scRNA-seq of FACS purified CD11b^+^ and CD11c^+^ (colon) or CD3/CD19/SiglecF/Ly6G^−^ CD11b^+^ (blood) from unmanipulated (naïve) C57BL/6J (WT) mice or WT mice fed 2% DSS for 6 days (acute colitis). **b.** Application of module score for scRNA-seq derived IAM signature using the top 20 specifically expressed genes from human *CXCL9/CXCL10* (Figure 1a) and mouse *Cxcl9/Cxcl10* monocytes (Figure 2a). **c.** Representative expression of CD319, CD274 and CCRL2 by colonic Ly6C^+^MHCII^−^ and Ly6C^+^MHCII^+^ monocytes from WT mice during acute colitis with 2% DSS (day 6). Data are representative of at least 10 independent experiments. **d.** Trajectory inference analysis derived by Monocle v3 from scRNA-seq analysis in a) using Pseudotime analysis (top) and cluster-diagrammatic overlay of major branches (bottom). **e.** Representative expression of tdTomato (representing active or historic expression of *Ms4a3*) by total Ly6C^+^ colonic monocytes or CD319-defined subsets obtained from acutely colitic (day 4) *Ms4a3*^Cre/+^.*Rosa26*^LSL-CAG-tdTomato/+^ reporter mice. Symbols represent individual mice. Data are from eight mice pooled from two independent experiments. Data analysed by paired student’s *t* test.

Monocyte heterogeneity of this kind has not been described previously in the inflamed intestine and while the accumulation of monocytes in the context of intestinal inflammation has been established by our previous work and that of others^15,16,23,32^, the appearance of a discrete monocyte cluster has not. To understand how IAM^mo^s relate to other monocyte clusters, we next performed trajectory-inference analysis on our paired blood-colon murine data. In support of existing work^10,33^, this analysis suggested that in steady state colon, *Ly6c2*^+^ blood monocytes were the precursors of colon generated *Cd163*^+^ and *Itgax*(CD11c)^+^ resident macrophages through a *H2-Aa*^+/−^ intermediary (**Figure 3a & d**). Whilst this observation also held in acute colitis, with the caveat that few mature macrophages were sampled, at the *H2-Aa*^+/−^ intermediary stage in acute colitis there was a branch-point to two discrete colon monocyte populations including IAM^mo^s, suggesting that this cluster may be unrelated, developmentally, to the CD319^−^ monocytic cells (**Figure 3d**). To test this directly, we used a genetic fate mapping model to track monocytes and their progeny. The *Ms4a3*^Cre/+^.*Rosa26*^LSL-CAG-tdTomato/+^ fate mapping model allows high levels of labelling of Ly6C^hi^ blood monocytes owing to their derivation from progenitors that transit through an *Ms4a3*^+^ state during their development^34^. If IAM^mo^s derived from CD319^−^ monocytes then they should label equally in this system. However, it was clear that IAM^mo^s were labelled at a significantly lower rate than their CD319^−^ counterparts (**Figure 3e**). Given the irreversible nature of labelling in this system, it is highly unlikely that IAM^mo^s derive from their CD319^−^ counterparts.

Taken together, these data support the premise that the transcriptional identity of IAM^mo^s is conferred locally after arrival in the inflamed intestine and they most likely represent a distinct lineage of monocytes that accrue in colitis.

### ACOD1 is an evolutionarily conserved feature of IAMs

To gain a better understanding IAM function, we next performed differential gene expression (DEG) analysis of IAM^mo^s in our combined naïve-acute colitis mouse *Csf1r^+^* dataset. This revealed 4,159 genes that were differentially expressed (2,312 higher and 1,847 lower) in IAM^mo^s compared with other *Csf1r^+^* cells (**Figure 4a**). The most upregulated gene was *Acod1*, which encodes the mitochondrial enzyme aconitate decarboxylase-1 (ACOD1; also known as immune responsive gene 1, IRG1) (**Figure 4a**). ACOD1 is the obligate source of the metabolite itaconate, which plays a number of anti-inflammatory roles^35^, but its effects on intestinal monocytes and macrophages are poorly understood. Importantly, *Acod1* was expressed exclusively by IAM^mo^s amongst *Csf1r^+^* cells (**Supp. Figure 4a**).

**Figure 4.**
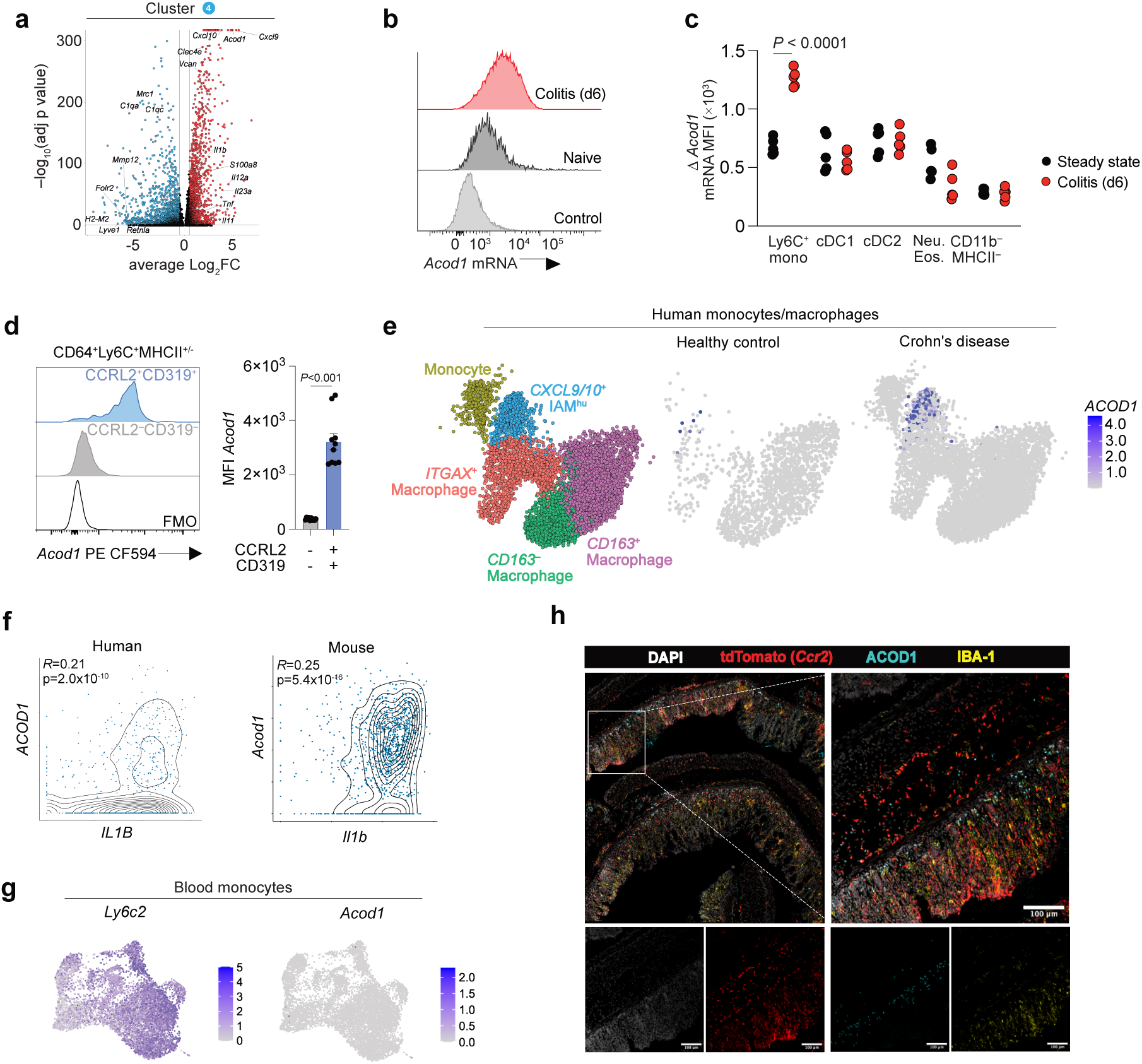
ACOD1 is an evolutionarily conserved feature of IAMs. **a**. Volcano plot of differential gene expression (DEG), with selected highlighted genes, comparing *Cxcl9/10^+^* cells with other *Csf1r^+^* cells in 2% DSS induced acute colitis (day6) from scRNA-seq in Figure 2a. **b-c**. Representative *Acod1* mRNA expression by colonic Ly6C^+^ monocytes (**b**) or indicated myeloid cell subsets (**c**) from naïve or 2% DSS treated C57BL/6J (WT) mice (day 6) as assessed by flow cytometry probe hybridisation (PrimeFlow). Symbols represent individual mice. Data from four mice per condition from one of three independent experiments, analysed by unpaired students *t* test. **d.** Representative *Acod1* mRNA expression by CD319 and CCRL2-defined Ly6C^+^ colonic monocytes from colitic (day 6) WT mice. Symbols represent individual mice. Data from ten mice pooled from three independent experiments, analysed by unpaired students *t* test. **e.** *ACOD1* expression in healthy human and active ileal CD colonoscopic biopsies as assessed by scRNA-seq data from Figure 1a. **f.** Scatter plot of *ACOD1/Acod1* and *IL1B/Il1b* by *CXCL9/10* and *Cxcl9/10* positive cells from Figure 1a and **2a**. analysed by Pearson correlation co-efficient. **g.** *Ly6c2* and *Acod1* expression by blood monocytes as assessed by scRNA-seq data from Figure 3a. **h.** Representative immunofluoresence of IBA1, ACOD1, DAPI and tdTomato (*Ccr2*) expression in distal colon sections of colitic (day 6) *Ccr2*^Cre-ERT2/+^.*Rosa26*^LSL-CAG-tdTomato/+^ mice administered tamoxifen at day 3.

To validate *Acod1* expression by a complementary method, we used PrimeFlow analysis, which measures mRNA expression at single cell resolution by flow cytometry. Consistent with our scRNA-seq observations, *Acod1* mRNA was negligibly present in all steady state cells examined. Again, consistent with our scRNA-seq observations, CD64^+^ Ly6C^+^ monocytes alone demonstrated large increases in *Acod1* expression in acute inflammation, where there was still little or no expression in any cDC subsets, granulocytes or other CD11b^−^ cells (**Figures 4b & c and Supp. Figure 4b**). Furthermore, within CD64^+^ Ly6C^+^ monocytes from acute colitis, it was CD319^+^ cells that were responsible for expression of *Acod1*, confirming IAM^mo^s as the major source of *Acod1* (**Figure 4d**). In keeping with this, our human dataset demonstrated that *ACOD1* expression was not found in any cell lineage other than IAM^hu^s, suggesting *Acod1/ACOD1* expression is IAM restricted (**Figure 4e**, **Supp. Figure 4c**). Finally, *ACOD1 / Acod1* levels were associated with the abundance of *IL1B / Il1b* transcript in IAMs (**Figure 4f**), implying that the highest IL-1β producing cells may also have the highest ACOD1 activity.

*Acod1* expression was also absent from blood monocytes, supporting its role as an exclusive feature of IAM education in the inflamed intestine (**Figure 4g**). Given their pro-inflammatory repertoire, we next hypothesised IAMs would localise in sites of high inflammatory burden. To address this, we localised IAM^mo^s in *Ccr2*^CreERT2/+^.*Rosa26*^CAG-LSL-tdTomato/+^ reporter mice in which CCR2^+^ cells, including classical monocytes and their immediate progeny, are labelled irreversibly with tdTomato in a tamoxifen-inducible manner. Administration of tamoxifen during DSS administration led to efficient labelling of cells in the colonic lamina propria and using immunohistochemistry, ACOD1^+^ cells could be clearly identified within the tdTomato^+^ fraction of cells. These were located preferentially in the regions of barrier damage, around the base of the crypts, consistent with recent tissue ingress (**Figure 4h**).

Taken together, these data show that IAM^mo^s accumulate near the sites of mucosal damage and are marked by high and unique expression of ACOD1 in mouse and human intestinal inflammation.

### Interferon and MyD88-dependent signals drive ACOD1 expression in IAMs

To understand the factors responsible for inducing IAM behaviour, we first performed gene set enrichment analysis (GSEA) of our human dataset, comparing IAM^hu^ DEGs in inflammation versus health. This revealed “IFN signalling” and “IFN-γ signalling” and “Interleukin signalling” as the most enriched pathways by IAM^hu^s (**Figure 5a**). Interestingly, using the same GSEA approach identified near identical top pathway enrichments in IAM^mo^s (“IFNα/β signalling”, “IFN-γ signalling” and “IL-1β signalling”) (**Figure 5b**), suggesting an evolutionary conserved response of these cells to intestinal damage.

**Figure 5.**
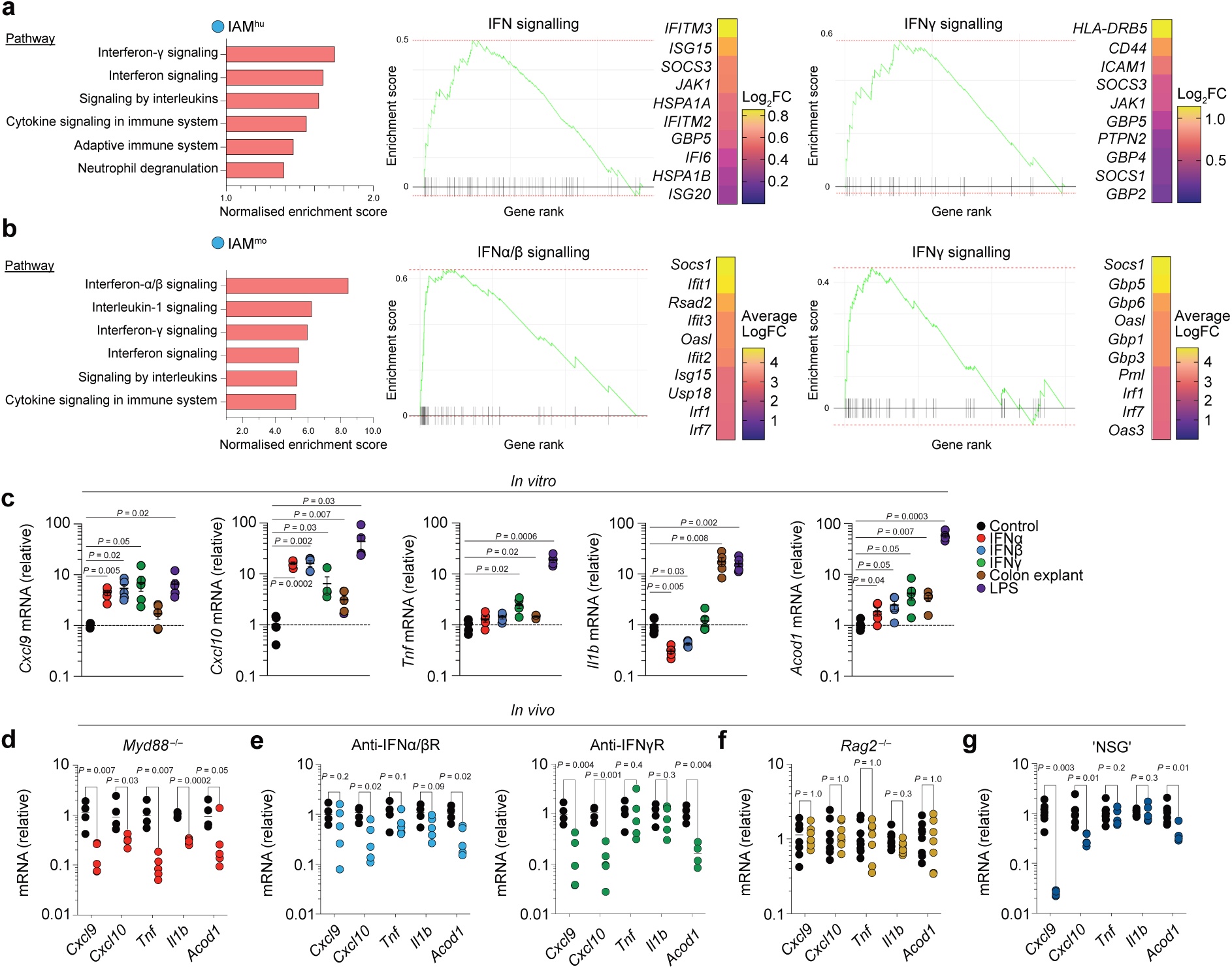
Interferon and MyD88-dependent signals drive ACOD1 expression in IAMs. **a-b**. Gene set enrichment analysis (GSEA) performed on **a)** *CXCL9/10*^+^ cells (IAM^hu^; Figure 1a) or **b)** *Cxcl9/10*^+^ cells (IAM^mo^; Figure 2a) with bar plots of top 20 pathways by normalised enrichment score (left) and enrichment plots (right) for IFNα/β and IFNγ signalling. Heatmaps show average expression of top enriched “leading edge” genes within each pathway. **c.** RT-PCR mRNA levels of indicated genes, normalised to *Actb,* by CD115^+^Ly6C^hi^ BM monocytes FACS purified from *Ms4a3*^Cre/+^*.Rosa26*^LSL-CAG-tdTomato/+^.*Cx3cr1*^+/gfp^ reporter mice, cultured with CSF-1 for 2days, then stimulated for 24 hours with rIFNα, rIFNβ, rIFNγ, LPS or the supernatant “secretome” from day6 DSS treated colon (colon explant). Symbols represent individual mice. Data are from five mice from one of two independent experiment performed, analysed by paired Students *t* test versus control. **d.** RT-PCR mRNA levels of indicated genes, normalised to *Actb,* by FACS purified Ly6C^+^ colonic monocytes from day 5, 2% DSS treated *Myd88^−/−^* mice relative to expression by FACS purified colonic Ly6C^+^ monocytes from colitic C57BL/6J mice. Symbols represent individual mice (n=4-5) from one experiment analysed by unpaired *t* test with multiple comparison (5% False discovery rate, two-stage step-up method). **e.** RT-PCR mRNA levels of indicated genes, normalised to *Actb,* by FACS purified Ly6C^+^ colonic monocytes from day 6, 2% DSS treated C57BL/6J mice treated with anti-IFNα/βR (clone MAR1-5A3) or anti-IFNγR1 (clone GR-20) or isotype control on day 2, 3, 4 and 5 of DSS administration. Symbols represent individual mice (n=4-5 from one experiment analysed by unpaired *t* test with multiple comparison (5% False discovery rate, two-stage step-up method). **f.** RT-PCR mRNA levels of indicated genes, normalised to *Actb,* by FACS purified Ly6C^+^ colonic monocytes from day 6, 2% DSS treated *Rag2^−/−^* mice relative to expression by FACS purified colonic Ly6C^+^ monocytes from colitic C57BL/6J mice. Symbols represent individual mice (n=7) pooled from two experiments, analysed by unpaired *t* test with multiple comparison (5% False discovery rate, two-stage step-up method). **g.** RT-PCR mRNA levels of indicated genes, normalised to *Actb,* by FACS purified Ly6C^+^ colonic monocytes from day 6, 2% DSS treated NSG mice relative to expression by FACS purified Ly6C^+^ colonic monocytes from colitic C57BL/6J mice. Symbols represent individual mice (n=5*) from two experiments analysed by unpaired *t* test with multiple comparison (5% False discovery rate, two-stage step-up method). *one sample was beneath the limit of detection for *Cxcl9*

IFN induction of *Acod1* expression in monocytes has been suggested in studies of mouse blood leukocytes stimulated with viruses, type I IFN or IFNγ *in vitro*^36–39^. However, as this work also found that various bacteria and TLR4 ligands could also induce *Acod1* expression, we examined how each of these pathways might contribute to the local induction of IAM^mo^s. First, we cultured Ly6C^+^ (CX3CR1^int^tdTomato^+^) monocytes purified from bone marrow (of *Ms4a3*^Cre/+^.*Rosa26*^CAG-LSL-tdTomato/+^.*Cx3cr1*^+/gfp^ reporter mice) with LPS, recombinant IFNα, IFNβ, IFNγ or the supernatant from inflamed colon explant tissue, “colon secretome”, and measured mRNA of five key IAM signature genes (**Figure 1b and 2d**) by RT-qPCR. LPS treatment led to significant upregulation of all signature genes examined (*Cxcl9, Cxcl10, Tnf* and *Il1b*), as well as causing profound upregulation of *Acod1* (**Figure 5c**). As expected, both type I and type II IFN led to upregulation of *Cxcl9* and *Cxcl10*, consistent with their IFN-inducible nature. However, expression of *Tnf* and *Il1b* was unaffected by type I IFNs, and IFNγ only elicited minor upregulation of *Tnf* in monocytes. Thus, full induction of the proinflammatory IAM signature involves the convergence of multiple signals. In contrast, *Acod1* expression was upregulated, albeit to different degrees, by all stimuli tested *in vitro*, including colon secretome (**Figure 5c**), demonstrating that the stimuli necessary for induction of ACOD1 expression are present in the inflamed colon.

To explore the induction of IAMs *in vivo*, we first used global *Myd88*^−/−^ mice in whom IL-1R and most TLR signalling is abrogated. Consistent with a non-redundant role in IAM^mo^ induction, purified CD64^+^Ly6C^+^ colonic monocytes from day6 DSS treated *Myd88*^−/−^ mice had significantly lower levels of mRNA for all genes within the five gene monocyte signature, including *Acod1,* compared with their WT counterparts (**Figure 5d**). Next, we abrogated type I or type II interferon pathways during DSS colitis by administering antibodies against either type I or type II IFN receptors (IFNAR1 and IFNγR1, respectively) and again isolated colonic monocytes for transcriptional profiling. Importantly, and consistent with our *in vitro* findings, blockade of either type I or type II IFN did not affect *Tnf* or *Il1b* expression in colonic monocytes, but reduced expression of *Cxcl10*, with anti-IFNγR but not anti-IFNAR1, also reducing *Cxcl9* expression (**Figure 5e**). Importantly and consistent with our *in vitro* data, *Acod1* expression was reduced significantly by both type I and type II IFN receptor blockade (**Figure 5d**).

To assess the relative contribution of lymphocytes and their products, which will include IFNs, to the monocyte signature, we first induced DSS colitis in *Rag2*^−/−^ mice and found that their colonic monocytes expressed the canonical IAM genes to the same extent as seen in wild type mice with colitis. Thus, mature T and B lymphocytes and/or their products are dispensable for IAM^mo^ induction in this model (**Figure 5f**). In contrast, monocytes from colitic ‘NSG’ (NOD *scid Il2rg*^−/−^) mice, which lack all mature lymphocytes including innate lymphoid cells (ILCs) and natural killer (NK) cells, had significantly lower expression of *Cxcl9, Cxcl10* and *Acod1,* whilst *Tnf* and *Il1b* were expressed at equivalent levels (**Figure 5g**). Together, these results suggest that although IFN and TLR signalling drive different proinflammatory facets of IAM induction, both are potent drivers of *Acod1* expression, with innate lymphocytes playing a crucial role in driving IAM^mo^ differentiation.

### ACOD1 regulates monocyte behaviour in inflammation

Given our previous data suggested *ACOD1/Acod1* is expressed exclusively in IAMs, we assessed the function of ACOD1 in IAM^mo^ *in vivo*, using *Acod1*^−/−^ mice^36^. Consistent with IAM specific expression, *Acod1* deficiency had no effect on the distribution of MDP, GMP and cMOP in BM (**Supp. Figure 5a**), steady state colon monocytes / macrophages (**Supp. Figure 5b**), or steady state colon tissue architecture (**Supp. Figure 5c**) with these parameters identical to those in WT controls. As the defining feature of IAMs was both high expression of pro-inflammatory mediators and ACOD1, the obligate source of the anti-inflammatory metabolite itaconate, we hypothesised that selective expression of ACOD1 in IAMs may represent an intrinsic mechanism to restrain excessive inflammatory behaviour.

To test this, we determined the impact of *Acod1* deficiency on the induction and recovery from DSS-induced colitis. In acute colitis, *Acod1*^−/−^ mice displayed increased weight loss compared with *Acod1*^+/+^ controls (**Figure 6a**), and flow cytometric analysis showed that this was accompanied by elevated numbers of Ly6C^+^ monocytes and CD11b^+^Ly6G^+^ neutrophils in the colon (**Figure 6b & c**). This was paralleled by significantly increased cellular infiltrates and colon shortening (**Supp. Figure 5d and e**). When we assessed the recovery from acute colitis, whereby mice are fed DSS for four days before returning to normal drinking water for up to 14 days (d4+14), we found that *Acod1*^+/+^ mice began to recover weight soon after DSS was withdrawn and all survived, whereas *Acod1*^−/−^ mice continued to show severe weight loss and ∼30% met protocol severity limits (**Figure 6d**). Furthermore, the surviving *Acod1*^−/−^ mice showed significantly heightened clinical scores compared with *Acod1*^+/+^ controls for almost 2 weeks after cessation of DSS (**Figure 6d**). Flow cytometric analysis at d4+14 showed that the *Acod1*^−/−^ mice also continued to have enhanced infiltration of the colon by CD64^lo/+^ Ly6C^+^ monocytes and CD11b^+^Ly6G^+^ neutrophils (**Figure 6e**) which was reflected in increased histopathology score and cellular infiltrate in *Acod1*^−/−^ mice (**Figure 6f & g**).

**Figure 6.**
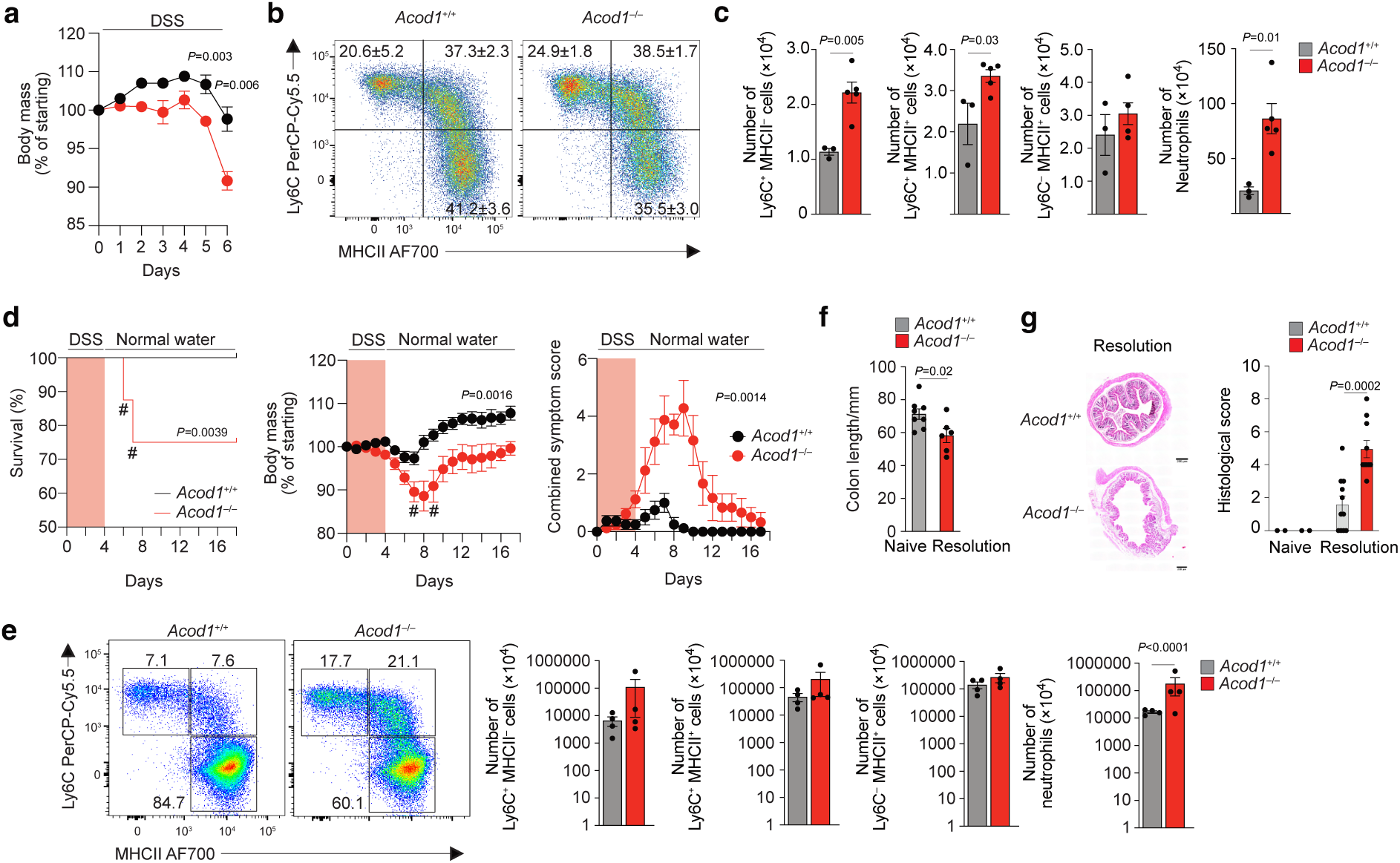
ACOD1 regulates monocyte behaviour in intestinal inflammation. **a.** Percentage body mass change compared to day0 starting weight of *Acod1*^+/+^ and *Acod1*^−/−^ mice over six days of DSS administration. Data from one of three experiments performed with n=5 mice per genotype, analysed by unpaired Students *t* test followed by a Holm-Śidák correction at each timepoint. **b.** Representative expression of Ly6C and MHCII by live CD45^+^CD11b^+^CD64^+^ cells from *Acod1*^+/+^ and *Acod1*^−/−^ mice at day 6 of DSS colitis. **c.** Absolute number of Ly6C- and MHCII-defined monocytes/macrophages and Ly6G^+^ neutrophils from *Acod1*^+/+^ and *Acod1*^−/−^ mice at day 6 of DSS colitis. Symbols represent individual mice (n=3-4) from one of two experiments performed, analysed by unpaired Students *t* test. **d.** Survival (left), body mass change (middle) and combined symptom score (right) of *Acod1*^+/+^ and *Acod1*^−/−^ mice fed 2% DSS for four days before being returned to normal drinking water, analysed by Log-rank (Mantel-Cox). # represents points where animals were culled due to reaching severity limits. **e.** Colon length from mice in **d**, analysed by unpaired Students *t* test. **f.** Histological score of H&E sections of distal colon from mice in **d**. Symbols represent individual mice (n=3-4) from one of three experiment performed, analysed by unpaired Students *t* test. **g.** Representative expression of Ly6C and MHCII by live CD45^+^CD11b^+^CD64^+^ cells from *Acod1*^+/+^ and *Acod1*^−/−^ mice at day 16 of the recovery model (left), representing twelve days post cessation of DSS. Graphs show absolute numbers of Ly6C- and MHCII-defined monocytes/macrophages and Ly6G^+^ neutrophils from *Acod1*^+/+^ and *Acod1*^−/−^ mice at day 16 of the recovery model of DSS colitis (middle-right). Symbols represent individual mice (n=3-4) from one of three experiment performed, analysed by unpaired Students *t* test.

Consistent with the role of ACOD1 in restraining pro-inflammatory cytokine production through the generation of itaconate^37,38^, the proportions of CD64^lo/+^Ly6C^+^ colonic monocytes from *Acod1*^−/−^ mice producing TNF and IL-1; were significantly increased compared with those from *Acod1*^+/+^ mice (**Figure 7a**), whereas deletion of *Acod1* did not alter the production of these cytokines by neutrophils (**Figure 7b**). To ascertain if supplementation with itaconate-derivatives would facilitate intestinal restitution during recovery from DSS colitis, C57BL/6J mice received 4-octyl itaconate (4OI), a cell permeable esterified-derivative of itaconate, intraperitoneally every 48hours in our resolution model of colitis (**Figure 7c**). Although this had no effect on the progress of disease as assessed by weight loss, 4’OI treated mice developed less diarrhoea (**Figure 7c**) and there were fewer colonic neutrophils and CD64^lo/+^Ly6C^+^ monocytes at d4+14 (i.e., the recovery period) **(Supp. Figure 5f, g and h).** However, the frequency of cytokine producing IAM^mo^s at this timepoint was equivalent in 4’OI treated and control mice (**Figure 7d & e**), suggesting that exogenous itaconate is insufficient to influence the proinflammatory behaviour of monocytes.

**Figure 7.**
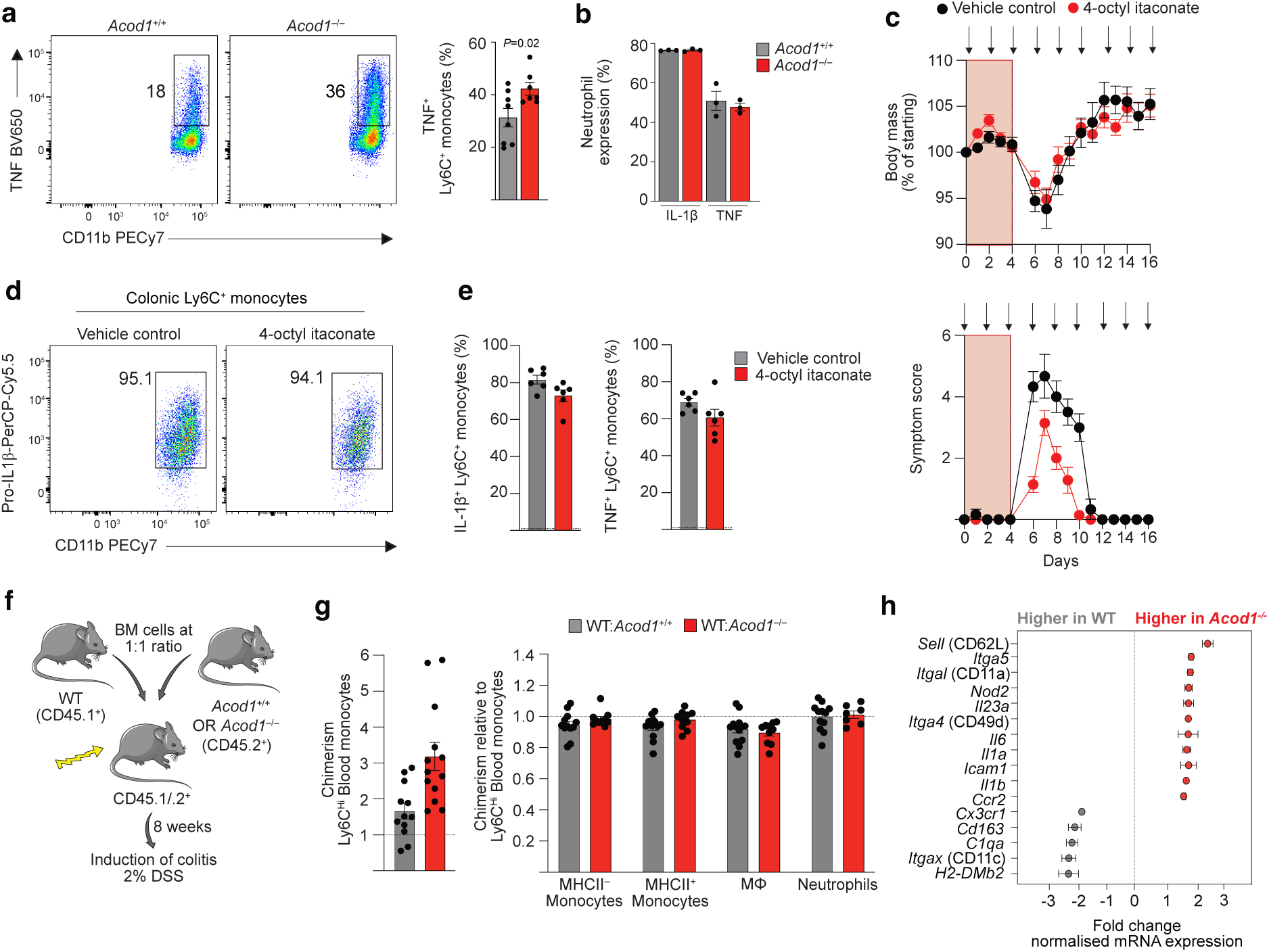
ACOD1 regulates cytokine responses in monocytes in a cell-intrinsic manner. **a.** Expression of TNF by CD64^+^Ly6C^+^ colon monocytes (after 3hours of *ex vivo* culture) as assessed by flow cytometry from *Acod1*^+/+^ and *Acod1*^−/−^ mice at day 16 of the recovery model, representing twelve days post cessation of DSS. Symbols represent individual mice (n=7-8) pooled from two independent experiments, analysed by unpaired Students *t* test. **b.** Frequency of TNF and IL-1β expressing colonic neutrophils in *Acod1*^+/+^ and *Acod1*^−/−^ mice as in **a**. **c.** Percentage body mass change of starting day 0 weight (top) and combined symptom score (bottom) of C57BL/6J mice fed 2% DSS for four days before being returned to normal drinking water and administered 4-octyl itaconate (4’OI) or vehicle control every 48hrs (indicated by arrows). Data from pooled from two independent experiments performed with n=5 (vehicle control) or n=8 (4’OI) mice per group. **d.** Expression of IL-1β by colonic Ly6C^+^ monocytes from mice in **c**. **e.** Frequency of IL-1β or TNF by colonic Ly6C^+^ monocytes from mice in **c**. **f.** Schematic of mixed bone marrow chimera generation. **g.** Chimerism of colonic Ly6C^+^ monocytes, Ly6C^−^MHCII^+^CD64^+^ macrophages and neutrophils in mixed WT:*Acod1*^−/−^ chimeric mice 8 weeks post reconstitution and after six days of DSS administration. **h.** Normalised expression of indicated genes by Ly6C^+^ monocytes, FACS purified from colitic WT:*Acod1*^−/−^ chimeric and analysed using nanoString nCounter analysis. Expression relative to that in WT (*Acod1* sufficient) cells in the same mouse with three biological replicates per genotype.

As ACOD1 expression is restricted to IAMs, these results suggest that deficiency of ACOD1 has a cell-intrinsic effect on monocytes. To examine this directly, we used a competitive BM chimera system in which lethally irradiated CD45.1^+^CD45.2^+^ mice were reconstituted with equal proportions of BM cells from *Acod1*^+/+^ (CD45.1^+^) and *Acod1*^−/−^ (CD45.2^+^), and acute DSS colitis was induced 8 weeks after reconstitution (**Figure 7f**). *Acod1*^−/−^ derived BM appeared to have a small selection advantage in blood chimerism (**Figure 7g**), but after normalising for this, *Acod1*^+/+^ and *Acod1*^−/−^ BM contributed equally to colon monocytes, macrophages and neutrophils (**Figure 7g**). Next, we FACS purified CD64^lo/+^Ly6C^+^ monocytes from these chimeric mice based on their expression of congenic CD45 expression and compared their transcriptome by nanoString technology (**Figure 7h**). This revealed that monocytes of *Acod1*^−/−^ origin had marked upregulation of genes pertaining to inflammatory responses (*Il1a, Il1b, Il6, Il23a* and *Nod2*) and recruitment (*Itgal, Itga5, Itga4* and *Icam*) (**Figure 7h**), whereas genes indicative of monocyte-to-macrophage maturation (*H2-Dmb2, Cx3cr1, C1qa, Itgax* and *Cd163*) were markedly downregulated (**Figure 7h**). As these cells come from an environment in which itaconate is available from *Acod1*-sufficient WT cells, these results demonstrate directly a cell-intrinsic role for ACOD1 in monocyte function.

Taken together, these results demonstrate that *Acod1* deficiency leads to excessive intestinal inflammation associated with the presence of dysregulated, hyperinflammatory monocytes and they suggest that ACOD1 acts as a cell-intrinsic molecular ‘hand brake’ to restrain IAM activity and differentiation.

## Discussion

Monocytes and macrophages are increasingly seen as central drivers of IBD^19^, yet also play a myriad of homeostatic roles to maintain gut health and orchestrate repair following injury^1^. How these cells can play such distinct roles in health versus inflammation remains poorly understood. We show that not all monocytes are equal, by identifying a distinct population of inflammation-associated monocytes (IAMs), which have superior pro-inflammatory functions and distinct developmental origin. Furthermore, we provide flow cytometry-based strategies to identify these cells without the need for advanced transcriptomics. We show the defining pro-inflammatory features of IAMs are imprinted by the inflamed intestinal environment, which also simultaneously triggers induction of an ACOD1 / itaconate-dependent regulatory mechanism. Thus, we show for the first time that dual pathways are selectively induced in IAMs to curtail pro-inflammatory function in human and murine intestinal inflammation.

The advent and application of technologies such as scRNA-seq has transformed our understanding of immune and non-immune cell heterogeneity, including in IBD^22,40,41^. While many of the published studies have documented (or ‘atlased’) heterogeneity across a variety of lineages^5,22,41–47^, few have specifically focussed on the changes within the monocyte / macrophage compartment. Indeed, Jahnsen and colleagues describe the only macrophage focused human single-cell study to date, using colorectal cancer resection specimens, where they identify 13 clusters of LP macrophage that could be broadly distinguished on the basis of *FOLR2/LYVE1, ACP5, CXCL2/3, S100A8/9* and *CXCL9/10*^29^.

However, those studies that include monocytes / macrophages in disease document marked changes to their heterogeneity and transcriptional state^5,22^. For example, Martin et al. describe a gene module derived from cell types that accumulate in CD, including ‘inflammatory macrophages’, that was associated with higher rates of failure to anti-TNF treatment. Our data complement and extend these findings. We now show additional heterogeneity can be found amongst monocytic cells than originally described in the first scRNA-seq study of CD^22^. Our data are also consistent with a study by Xavier and colleagues ^5^ which also identified marked expansion of monocytic cells in CD patients, with *S100A8/9^+^*and *CXCL9/CXCL10^+^* subsets as we identified here, although in their data *CXCL9/CXCL10^+^* cells did not show significant expansion in non-inflamed versus inflamed ileal tissue. Moreover, they also found highest enrichment for IBD risk genes *NOD2, LRRK2, JAK2, TYK2, IFIH1* and *NFKB1* in monocytic subsets. Our analysis also identified surface markers for the identification of IAMs, namely CD319, CD274 and CCRL2. *SLAMF7,* which encodes CD319, has previously been identified as feature of highly inflammatory cells and has been proposed to potentiate proinflammatory responses through homotypic interactions. While none of these surface proteins can be used as *de facto* markers of IAM, their expression in combination with canonical monocytic markers, such as CD14, CD55 and CD11a, allows IAMs to be identified. That *SLAMF7* has been shown to mark proinflammatory macrophages in the context of rheumatoid arthritis and severe COVID-19 disease, suggests this is a general feature of proinflammatory monocytes and macrophages rather than specific to IBD.

Our scRNA-seq approach also allowed putative relationships between IAMs and other monocyte subsets to be examined. Our results suggested that distinct trajectories exist within the monocyte compartment in the intestinal mucosa, intriguing considering recent work describing heterogeneity in blood monocytes^48,49^. Specifically, it has been proposed that monocytes can derive from either monocyte dendritic progenitors (MDPs) or granulocyte macrophage progenitors (GMPs) with a relative contribution to the circulating pool dependent on the environmental context^48,49^. MDP- and GMP-derived cells are differentially labelled in the *Ms4a3*^Cre/+^.*Rosa26*^LSL-CAG-tdTomato/+^ fate mapping model we used here, with higher labelling in GMP-derived monocytes^34^. Moreover, MDP-derived monocytes have been suggested to be enriched for *Slamf7/*CD319 expression in blood^49^. Thus, given we noted reduced genetic labelling in IAM^mo^s (in colitic *Ms4a3*^Cre/+^.*Rosa26*^LSL-CAG-tdTomato/+^ mice), it is plausible that IAMs may represent MDP-derived cells rather than conventional GMP-derived cells, although establishing the exact ontogeny of intestinal monocyte / macrophage subsets was not the focus of this study and warrants further investigation with new, specific lineage tracing tools.

Irrespective of developmental origin, our paired scRNA-seq analysis (blood-colon) supports the premise that the signature of IAMs is primarily driven by environmental factors in the intestinal mucosa. In both our human and mouse data, expression of ACOD1 was one of the most specific features of IAMs. Indeed, upregulation of ACOD1 is well established in response to different proinflammatory stimuli, although much of this work is based on *in vitro* systems. Our work shows that ACOD1 induction is highly specific to IAMs and does not occur in other monocyte or macrophage subsets in the same mucosal biopsy. This likely reflects concentration of IAMs in high-damage / inflamed areas, which is supported by a recent study using spatial transcriptomics in active IBD whereby a *CXCL5* and *ACOD1* expressing macrophage subset was shown to preferentially locate at inflamed areas ^50^. Moreover, CD14^+^CXCL9^+^ monocytic cells, which are likely to similar to IAMs, have been shown to be enriched in areas of damage and neutrophil accumulation^51^.

The prominence of *CXCL9* and *CXCL10* within the IAM signature implicate IFNs in their generation. Indeed, blockade of either type I or type II IFN receptor signalling *in vivo* diminished the expression of these features, although not of effector cytokines *Tnf* or *Il1b*, which could be induced by LPS. Our results shows that the full IAM signature is dependent on the convergence of multiple signals. Importantly, ACOD1 was induced by all proinflammatory stimuli tested *in vitro*, findings consistent a number of previous studies^36,36,38^. Little work has interrogated ACOD1-inducing factors *in vivo* in the context of tissue inflammation. Our work provides *in vivo* evidence that TLR ligands and IFNs are potent inducers of ACOD1, although we do not dispute that other untested factors may also induce ACOD1. While our experiments in mice suggest that NK cells and ILCs (non-RAG-dependent lymphocytes) are indispensable for induction of the IAM signature, and ACOD1 in particular, our mouse colitis model does not replicate the chronicity of IBD where T cells will play a more prominent role. Indeed, CD14^+^CXCL9^+^ monocytic cells have been shown to be enriched in areas with high numbers of *IFNG*^hi^ CD8^+^ T cells in acute IBD^51^. Thus, it is likely that in IBD various sources contribute to the stimuli triggering ACOD1 expression.

Our results showing exacerbated intestinal inflammation in the context of *Acod1* deficiency supports the well-documented role of ACOD1-dependent itaconate as an anti-inflammatory mechanism within myeloid cells^52^. While the increased susceptibility of *Acod1* deficient mice to DSS colitis has been reported before^53^, the cellular basis for this has not been examined. We demonstrate that excessive monocyte pro-inflammatory behaviour is a feature of the heightened inflammation in *Acod1*^−/−^ mice. Although we used global *Acod1*^−/−^ mice and therefore cellular dysfunction caused by *Acod1* deficiency could plausibly be in any cell lineage, the very specific expression of ACOD1 within IAMs strongly suggests the primary dysfunction lies within these cells. There are a variety of mechanisms by which itaconate can elicit anti-inflammatory effects, including inhibiting succinate dehydrogenase (SDH), thereby increasing intracellular succinate, which regulates HIF-1α, decreasing IL-1β and ROS production. Itaconate can also activate NRF2 (nuclear factor erythroid 2-related factor 2) and enhance ATF3, both of which inhibit the inflammasome component, NLRP3^39^. Itaconate is also reported to have direct anti-microbial function, in part through inhibition of the glyoxylate cycle enzyme isocitrate lyase (ICL)^38,54^. ICL blockade restrains microbial energy production in low glucose environments thus limiting exponential growth phase in acute infection. However, the concentrations of itaconate required to produce these effects may be many times greater than those reported in macrophages *in vivo*, though this may be overcome by concentration in intra-cellular vacuoles^38,54^.

Importantly, we identify that IAM-intrinsic ACOD1 is vital for appropriate monocyte function, as exogenous itaconate by synthetic itaconate-derivative supplementation is not sufficient to overcome constitutive lack of *Acod1.* While these results could reflect an inability of certain exogenous itaconate-derivatives to enter cells effectively, as has been reported using *in vitro* assays where supra-physiological concentrations are needed to elicit effects ^55^, our data using competitive chimeric mice determined that cell intrinsic itaconate is indispensable for regulation of monocyte function. Our results are consistent with a very recent study where THP1 monocytic cells were used to demonstrate the importance of cell-intrinsic itaconate in controlling monocyte / macrophage responses^55^. However, this study was conducted entirely *in vitro*, thus to the best of our knowledge, our study represents the first demonstration of this function *in vivo*.

Taken together, these data suggest that the intestinal micro-environment directs a hyper-inflammatory subset of monocyte to areas of barrier damage. Simultaneously, the intestine also instructs the first phase of inflammation resolution, by initiating an immunometabolic programme exclusively within these cells, under the control of ACOD1. These data suggest mechanisms to augment intra-cellular itaconate may be of therapeutic benefit in pathogenic inflammatory states, such as IBD.

## Materials and Methods

### Mice

C57BL/6J and C57BL/6N mice were obtained from Charles River Laboratories, C57BL/6N-*Acod1*^em1(IMPC)J^/J (referred to as *Acod1*^−/−^) were obtained originally from Jackson Laboratories, but were provided by Prof. A Byrne (Imperial College, London), B6.SJL-*Ptprc*^a^*Pepc*^b^/BoyJ (referred to as CD45.1^+^) and B6.SJL-*Ptprc*^a^*Ptprc*^b^*Pepc*^b^/BoyJ (referred to as CD45.1/.2^+^) were bred in house at University of Edinburgh, *Ccr2*^Cre-ERT2-mKate^ were obtained from Prof. B. Becher (University of Zurich)^56^, C57BL/6J-*Ms4a3*^em2(cre)Fgnx^/J (referred to as *Ms4a3*^Cre^) were obtained from Prof. Florent Ginhoux (Singapore Immunology Network, {SIgN, A*}), B6.Cg-*Gt(Rosa)26Sor*^tm14(CAG-tdTomato)Hze^/J (referred to as *Rosa26*^LSL-CAG-tdTomato^) were obtained from Dr. Rebecca Gentek (University of Edinburgh), B6.Cg-*Rag2*^tm1.1Cgn^/J (referred to as *Rag2*^−/−^) were obtained from Prof. Anna Williams (University of Edinburgh), B6.129P2(SJL)-*Myd88*^tm1.1Defr^/J (referred to as *Myd88*^−/−^) were obtained from Prof. Owen Sansom (CRUK Scotland Institute), NOD.Cg-*Prkdc*^scid^*Il2rg*^tm1Wji^/SzJ (referred to as ‘NSG’ mice) were obtained from Charles River Laboratories. All mice were housed under specific pathogen-free conditions at the University of Edinburgh on a 12hr light/12hr dark cycle (light from 07:00-19:00 and dark from 19:00-07:00) at 22°C and 42% humidity. Mice were fed standard chow *ad libitum*. All experimental mice were age matched, with littermate controls where possible, and both sexes were used throughout the study. The mice used in each experiment is documented in the appropriate figure legend. Experiments performed at UK establishments were permitted under license by the UK Home Office and were approved by the University of Edinburgh Animal Welfare and Ethical Review Body (AWERB). Genotyping was performed by Transnetyx using real-time PCR.

### Human subjects

The acquisition of samples for human scRNA-seq was approved by the National Health Service (NHS) Research Ethics Committee (Cambridge South, REC ID 17/EE/0338). Written informed consent was given by all participants. Individuals undergoing routine endoscopic assessment were recruited at Addenbrooke’s hospital, Cambridge, UK. All CD participants had a confirmed history of CD and macroscopic evidence of terminal ileal inflammation from tissue sampled during the biopsy. All control participants were undergoing endoscopic assessment or surveillance for healthy and non-cancer related reasons (e.g., history of iron deficiency anaemia, family history of colorectal cancer). Control participants did not have macroscopic evidence of intestinal inflammation, a personal history of cancer, and were not in receipt of corticosteroids or any other immune modulating therapy. Patients who were taking probiotics or antibiotics were excluded. Patients of non-European ancestries were also excluded to reduce confounding. Pinch-biopsies of the terminal ileum were collected from all participants and deposited into pre-chilled Hanks Balanced Salt Solution (HBSS) without Mg^2+^, Ca^2+^, or phenol red. Samples were placed on ice and immediately transferred to the Sanger Institute.

For flow cytometry validation experiments, biopsy samples were obtained from adult IBD (both CD and UC) patients or healthy controls attending routine colonoscopy for initial disease surveillance or ongoing disease assessment (see **Supp. Table S2** for anonymized patient information) at the Western General Hospital, Edinburgh, UK, after informed consent under existing approvals (REC:19/ES/0087). A diagnosis of IBD was (based on standard clinical, endoscopic, radiological, and histological criteria). Endoscopic assessment of disease severity at the biopsy site was performed by clinicians using the Mayo endoscopic sub-score for UC or the simple endoscopic score (SES-CD) for CD and classified as quiescent, mild, moderate or severe. Two to four biopsies were taken per site and pooled for analysis.

### Tamoxifen administration

For induction of Cre activity in *Ccr2*^Cre-ERT2/+^.*Rosa26*^LSL-CAG-tdTomato/+^ mice, tamoxifen (T5648-1G, Scientific Laboratory Supplies) was dissolved in sesame oil overnight at 50mg/ml in a glass vial and administered by oral gavage at 5mg per mouse at day 3.

### In vivo IFN receptor blockade

Anti-mouse IFNgR1 (clone GR-20), anti-mouse IFNAR1 (clone MAR1-5A3) and mouse IgG1 isotype control were obtained from BioXcell, diluted in sterile PBS and administered at 250mg/mL intraperitoneally per mouse per day on days 2, 3, 4 and 5 of acute DSS colitis. Analysis of FACS-purified colonic monocytes was assessed by RT-qPCR on day 6 of colitis.

### 4-octyl itaconate administration

4-octyl itaconate (25374-100mg-CAY) was prepared fresh in 85% sterile water, 10% CyclodextrinB (Sigma-Aldrich) and 5% KollihorHS15 (Sigma-Aldrich) at a concentration of 5mg/ml. Mice received 50mg/kg by intraperitoneal injection every 48hours for 14days.

### DSS colitis

To induce colitis, mice received 2% dextran sodium sulphate (DSS) (reagent grade; MW 36,000-50,000 kDa; MP Biomedicals, Ohio) *ad libitum* in sterile drinking water for up to six days (acute colitis) or for four days before being returned to normal drinking water (recovery model). Mice were scored daily for weight loss and symptom score. Specifically, we quantified stool consistency: Well-formed/normal = 0, Pasty/semi-formed = 1, Pasty = 2, Diarrhoea that does not adhere to anus = 3, Diarrhoea that adheres to anus = 4, Weight loss: 1-5% = 1, 5-10% = 2, 10-20% = 3, ≥20% =4 and General appearance: Piloerection = 1, lethargy and piloerection = 2, sickly sunken eyes, ataxic = 4.

### Generation of competitive bone marrow chimeric mice

8-10 week old CD45.1^+^CD45.2^+^ WT mice were lethally irradiated with two doses of 4.6 Gy 3 hours apart before being reconstituted within 3 hours with 5×10^6^ WT (CD45.1^+^) and *Acod1*^−/−^ (CD45.2^+^) bone marrow cells at a ratio of 1:1. Controls received mixed bone marrow from WT (CD45.1^+^) and *Acod1*^+/+^ (CD45.2^+^). Chimerism was assessed pre-reconstitution and 8 weeks post-reconstitution by analysis of blood leukocytes by flow cytometry.

### Isolation of murine bone marrow, blood and intestinal leukocytes

Mice were sacrificed by cervical dislocation or by overdose with sodium pentobarbitone. Colonic leukocytes were isolated as described previously^10^. Briefly, the large intestine was excised, measured and soaked in PBS before any fat tissue was removed. If collecting tissue for histological analysis, a 0.5cm section of distal colon was placed in AntigenFix (Diapath) or formalin. Colonic tissue was incubated twice, each with 5mL 2mM EDTA/HBSS at 37°C for 15mins, in a shaking incubator before digestion in 10%FCS/RPMI1640 with 0.625mg ml*^−^*^1^ collagenase D (Roche), 0.425mg ml*^−^*^1^ collagenase V (Sigma-Aldrich), 1mg ml*^−^*^1^ Dispase (ThermoFisher), and 30 U ml*^−^*^1^ DNase (Roche Diagnostics GmbH) for 35 minutes in a shaking incubator at 37°C. Cell suspensions were washed in FACS buffer (2% FCS/2mM EDTA/PBS) and kept on ice until staining for flow cytometry. To isolate bone marrow cells, tibiae and femora from mice were excised, all flesh removed and washed in 70% ethanol. The ends of the bones were cut off and bone marrow flushed with sterile PBS using a 25G needle into a Petri dish.

### Human intestinal leukocyte isolation

For human scRNA-seq analysis, terminal ileal biopsies were dissociated using a single-step digestion protocol on ice to release all major intestinal cell types present in the biopsy (epithelial, immune, and stromal) without stressing the cells. First, the biopsies were mechanically minced and pipetted to release immune cells (fraction 1) from the lamina propria and the remaining tissue chunks were transferred to HBSS^−/−^ containing 2 mM EDTA, 0.26 U/µl serine endoprotease isolated from *Bacillus licheniformus* (Sigma, P5380), 5 µM QVD-OPh (Abcam, ab141421), and 50 µM Y-27632 dihydrochloride (Abcam, ab120129). Tissue chunks were pipetted regularly during a 30-minute incubation on ice to release epithelial and stromal cells (fraction 2). The cells from both fractions are washed, centrifuged, and then incubated for 10 minutes at room temperature in Hank’s Balanced Salt Solution (HBSS) with Mg^2+^, Ca^2+^, and without phenol red, including 5 mM CaCl_2_, 1.5U/µl collagenase IV (Worthington, LS004188), and 0.1 mg/ml DNase I (Stem Cell Technologies, 07900). The cells were then filtered (30 µm; CellTrics 04-0042-2316), washed, and centrifuged before being incubated for 3 minutes at room temperature in the red blood cell lysis buffer (ACK lysis buffer; Gibco, A10492).

For flow cytometry analysis, colonoscopic biopsy samples were collected in ice cold HBSS/2% FCS. Tissue was then placed in 5mls RPMI/5%FCS/1%PenStrep (“R5”) containing 4mM DTT for 10mins in a shaking incubator at 37°C, 180RPM. Samples were then washed in R5 before being placed in HBSS(CMF)/5mM EDTA for 10mins, washed in R5, followed by a further 10min HBSS(CMF)/5mM EDTA incubation, in a shaking incubator at 37°C, 180RPM. Samples were washed in R5, then digested in R5 containing 1.2mg/ml Liberase TM (Roche) for 30mins in a shaking incubator at 37°C for 30mins, 200RPM. Samples were then dissociated with a Pasteur pipette (30times), diluted with 30mls ice cold R5, then passed through a 100μm strainer before centrifugation at 400g for 5mins.

### Flow cytometry

For analysis of unfixed cells, cells were first incubated with 0.025 μg anti-CD16/32 (2.4G2; Biolegend) for 10mins on ice to reduce non-specific binding to Fc receptors and then stained with a combination of the antibodies detailed in **Supp. Table S3**. Where appropriate, cells were then stained with streptavidin-conjugated BV650 (Biolegend). Dead cells were excluded using DAPI or 7-AAD (Biolegend) added 2mins before acquisition. When assessing intracellular markers, cells were first washed in PBS and then incubated with Zombie NIR fixable viability dye (Biolegend) for 10mins at room temperature protected from light. Following the final wash step, cells were fixed and permeabilized using FoxP3/Transcription Factor Staining Buffer Set (eBioscience), and intracellular staining performed using antibodies detailed in **Supp. Table S3**. Samples were acquired using an LSRFortessa using FACSDiva software (BD) and analyzed with FlowJo software (version 9 or 10; BD). Analysis was performed on single live cells determined using forward scatter height (FCS-H) versus area (FSC-A) and negativity for viability dyes. mRNA was detected by flow cytometry using PrimeFlow technology (ThermoFisher) using probes against *Acod1* (AF647) according to the manufacturer’s guidelines. For staining controls in PrimeFlow analysis, the Target Probe Hybridization step was omitted. For detection of intracellular cytokines, whole colonic isolates were incubated in complete RMPI together with monensin 1μl/ml (GolgiStop, BD) for 3 hours at 37°C, before being washed and stained as above.

### In vitro analysis of monocytes, MDP and GMP

Bone marrow was isolated *Ms4a3*^Cre/+^.*Rosa26*^CAG-LSL-tdTomato/+^ reporter mice and MDP (CD117^hi^CD16/32^−^CD135^+^CD34^+^CD115^+^Ly6C^−^), GMP (CD117^+^CD16/32^+^CD34^+^CD115^−^ Ly6C^−^) and Ly6C^+^ monocytes (CD117^−^CD16/32^+^CD34^−^CD115^+^Ly6C^+^) were purified by FACS. MDP and GMP were then cultured for up to three days with recombinant 20ng/mL CSF1 (Peprotech) before analysis by flow cytometry for tdTomato expression. Monocytes were cultured for three days with CSF1 prior to addition of recombinant IFNα (100ng/ml), IFNβ (20ng/ml), IFNγ (80ng/ml), LPS (100ng/ml) derived from *E.coli* or supernatant from explanted inflamed colonic tissue (100AU/ml). After which cells were harvested and RNA extracted for transcriptional analysis.

### Immunofluorescence

Colons from day 6 DSS treated *Ccr2*^Cre-ERT2/+^.*Rosa26*^LSL-CAG-tdTomato/+^ mice were opened longitudinally and washed with ice cold PBS. Colons were then rolled distal-proximal (“Swiss-roll”) and placed in AntigenFix (Diapath) for 5mins, before washing in PBS and then 33% PBS/sucrose for 23hours. Samples were then snap frozen on a dry ice-methanol ice bath in OCT (Leica), with Swiss rolls being cryosectioned (Leica) at 20μm thickness. Prior to staining, sections were treated with methanol-acetone for 1min and then air-dried. Sections were then rehydrated in PBS for 5mins, blocked for 30mins (2%BSA, 1%Donkey serum 0.5% saponin), washed in PBS and then stained with primary antibodies overnight at 4°C (See **Supp. Table S3).** For ACOD1 staining, an Akoya OPAL polymer HRP kit was used before secondary antibody staining, isotype control, and DAPI. Slides were then fixed in ProlongGold and imaged by confocal microscopy.

### Histology

1cm distal sections of colon were placed in 4% paraformaldehyde for 24hours, before being placed in 70% Ethanol, cut into 20μm sections and stained with H&E. Slides were imaged using an AxiosSlideScanner and histological assessment of colitis was performed blinded using an established scoring system^57^. Each colonic sample was graded semi-quantitatively (Scores 0-3 per component): A. Degree of epithelial hyperplasia and goblet cell depletion. B. Leukocyte infiltration in the lamina propria. C. Area of tissue affected. D. Presence of markers of severe inflammation (such as crypt abscesses, submucosal inflammation and presence of ulcers).

### Human scRNA-seq

Single-cell RNA sequencing was undertaken using 3’ 10X Genomics kits (v3.0 and v3.1) according to the manufacturer’s instructions. All samples sequenced under kit version v3.1 had dual indexes, samples sequenced under kit version v3.0 had either single or dual indexes. Libraries were sequenced using a HiSeq4000 sequencer or NovaSeq S4 XP sequencer with 100bp paired-end reads, targeting 50,000 reads per cell. CellRanger v3.0.2 was used to demultiplex reads, align reads to GRCh38 with Ensembl version 93 transcript definitions (GRCh38-3.0.0 reference file distributed by 10X Genomics), and generate cell by gene count matrices. CellBender v2.1 was then applied to identify droplets containing cells and adjust the raw counts matrix for background ambient transcript contamination^58^. The final counts matrix was adjusted for the ambient transcript signature at a false positive rate of 0.1. Next, multiplets were identified and removed (scrublet v0.2.1)^59^.

### De novo cell type identification

The atlasing cohort (26 Crohn’s disease and 25 healthy participants) was used to identify cell types and fit a model to automatically predict cell types across the entire 121 subject dataset. Subsequent processing and management of the expression data was performed using scanpy v1.6.0^60^. After mean centering and scaling the ln(CP10K+1) expression matrix to unit variance, principal component analysis (PCA; sc.tl.pca) was undertaken using the 2,000 most variable genes after removal of protein coding mitochondrial, ribosomal, and immunoglobulin genes, because these genes constituted the ambient signature learned by CellBender. Clusters were defined using the Leiden graph-based clustering algorithm v0.8.3 on the nearest neighbours determined by bbknn. Clusters were generated across a range of resolutions from 0.5 to 5 to empirically determine the optimal clustering resolution. For each resolution considered, the data was divided into training (2/3 of cells) and test (1/3 of cells) sets and a single layer dense neural network fit to predict cluster identity from expression using keras v2.4.3. For extended methods and non-myeloid data, please see our pre-print^26^.

### Mouse scRNA-seq

Female C57BL/6J mice were treated with 2% DSS in drinking water for 6days or normal drinking water control (n=5 per group), and at day6. Myeloid cells (Live/CD45^+^/CD3,CD9^−^/CD11b^+^ or CD11c^+^ were enriched using a BD Fusion cell sorter and collected into PBS containing 1% BSA. Cells were checked for number and viability with 1:1 Trypan Blue under a brightfield microscope. This was performed separately for samples collected from each individual mouse. Following this, samples for each experimental group were mixed together to create a mixture of cells from two individual biological replicates with a final total of 20,000 cells. The cells were processed for single cell barcoding using the 10x Chromium platform and a library prepared for each sample using the 10X Genomics single-cell RNA-seq 3’ V3.1 kit. Sequencing was performed on the NovaSeq 2×150bp platform to a depth of 50-75,000 reads per cell.

### Pre-processing of murine scRNA-seq data

Pre-processing of the data was performed using Seurat v5.0 according to the workflow suggested by the Satija lab^61^. First, ambient RNA was identified by comparing the raw and filtered matrices and contaminant RNA estimated using the SoupX package. Adjusted matrices were then imputed into the R environment (v3.4) and analysed using Seurat. Normalisation was performed using regularised negative binomial regression via the SCTransform function, including the removal of confounding variation arising from mitochondrial mapping percentage. Doublets were identified and removed using the “Doublet Finder” package via artificial next nearest neighbour analysis. Batch effect correction was assessed using the Harmony package. Identification of highly variable genes, next nearest neighbour, clustering functions and UMAP visualisation were all performed using the Seurat package. Marker genes per identified subpopulation were found using the findMarker function of the Seurat pipeline. For trajectory inference analysis, Monocle 3 was used^62^.

### Nanostring

For murine colon monocytes isolated from day6 2% DSS treated CD45.1/2 chimeric mice, 25,000 CD45.1 (WT) and CD45.2 (*Acod1^−/−^*) CD64^+^Ly6C^+^ cells were isolated by FACS directly into 200 μl of RLT buffer using a BD Fusion Sorter from 5 mice. Cell pellets were vortexed and centrifuged before immediate freezing until ready for processing. NanoString gene expression plates using mouse immunology v2 panels were run as per the manufacturer’s instructions at the University of Edinburgh HTPU Centre within the MRC Institute of Genetics and Molecular Medicine/Cancer Research UK Edinburgh Centre.

### Quantitative PCR analysis (RT-qPCR)

RNA was isolated from whole tissue or sorted cells using the RNAqueous Micro Kit (Life Technologies). Total RNA samples were DNase-treated (Life Technologies) prior to cDNA synthesis (First Strand Synthesis Kit, ThermoFisher). SYBRgreen qPCR was performed using 5ng cDNA and primers designed using Primer3 and Beacon Designer software or described on PrimerBank. Primer sequences are listed in **Supp. Table S4**. Melt-curves and primer efficiency were determined as previously described. Gene expression was normalized to the housekeeping gene *Actb* and to the corresponding experimental control. Reactions were run in duplicate.

### Statistical analysis

Statistics were performed using Prism 10 (GraphPad Software). The statistical test used in each experiment is detailed in the relevant figure legend

## Acknowledgements

We are grateful to Prof. Florent Ginhoux for the provision of *Ms4a3*^Cre^ mice, to Prof. Burkard Becher for providing the *Ccr2*^Cre-ERT2-mKate^ mice, to Prof. Owen Sansom and Dr. Ed Roberts for providing the *Myd88*^−/−^ mice, to Dr. Rebecca Gentek for providing the *Rosa26*^LSL-CAG-tdTomato^ mice and to Prof. Anna Williams for providing the *Rag2*^−/−^mice. Furthermore, we would like to thank Rachel Ridgway for performing DSS in Myd88 deficient mice. Flow cytometry data were generated with the support from the IRR Flow Cytometry and Cell Sorting Facility, University of Edinburgh. We would like to thank Mariana Beltran and Elena Sutherland for technical expertise in setting up 10X sequencing and to Dr John Wilson-Kanamori for initial processing of the scRNA-seq raw data. We would also like to thank Prof. Prakash Ramachandran and Dr. Bert Malengi for their advice in scRNA-seq analysis, Ms Alison Munro for her expertise in Nanostring nCounter processing and Dr. Runchi Bansal for her advice on 4-octyl-itaconate administration. Finally, we would like to thanks the Bioresearch and Veterinary Services team at the University of Edinburgh for husbandry of our mice and other technical assistance.

## Author Contributions

G.R.J. conceived, performed, analysed experiments and wrote the manuscript. B.D. and L.H. performed and analysed experiments. T.A. and M.K. performed human scRNA-seq analysis. A.B. advised on *Acod1^−/−^* experiments and edited the manuscript. C.A. supervises T.A. and M.K. and edited the manuscript. T.R. assisted with human scRNA-seq analysis, interpretation, obtained tissue samples and edited the manuscript. G.T.H. obtained tissue samples, edited the manuscript and co-supervised the project. C.C.B. conceived, performed and analysed experiments, co-wrote the manuscript and supervised the project. All authors had the opportunity to edit the manuscript.

## Data and material availability

All data needed to evaluate conclusions in the paper are present in the paper or Supplementary Materials. The latest human scRNA-seq data can be accessed at (Zendo, DOI:10.5281/zenodo.8301000), mouse scRNA-seq data will be deposited in the National Center for Biotechnology Information Fene Expression Omnibus public database (www.ncbi.nlm.nih.gov.geo/).

