## Supplementary Material for "Inflammation-associated monocytes express ACOD1 to curtail inflammatory behaviour in IBD and experimental colitis"

### SUPPLEMENTARY MATERIAL Jones et al.

#### Figure S1

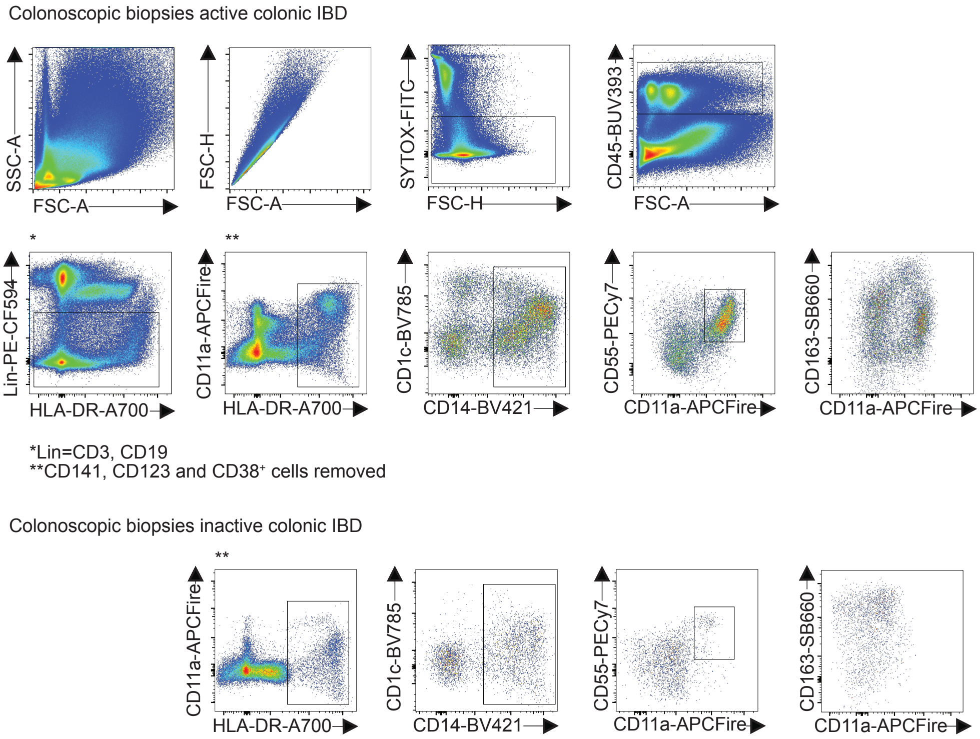

**Figure S1 Human IBD flow cytometry macrophage identification strategy.**
Representative flow cytometry analysis of mucosal biopsies from macroscopically inflamed (active) or uninflamed (inactive) IBD patients obtained at colonoscopy.

### Figure S2

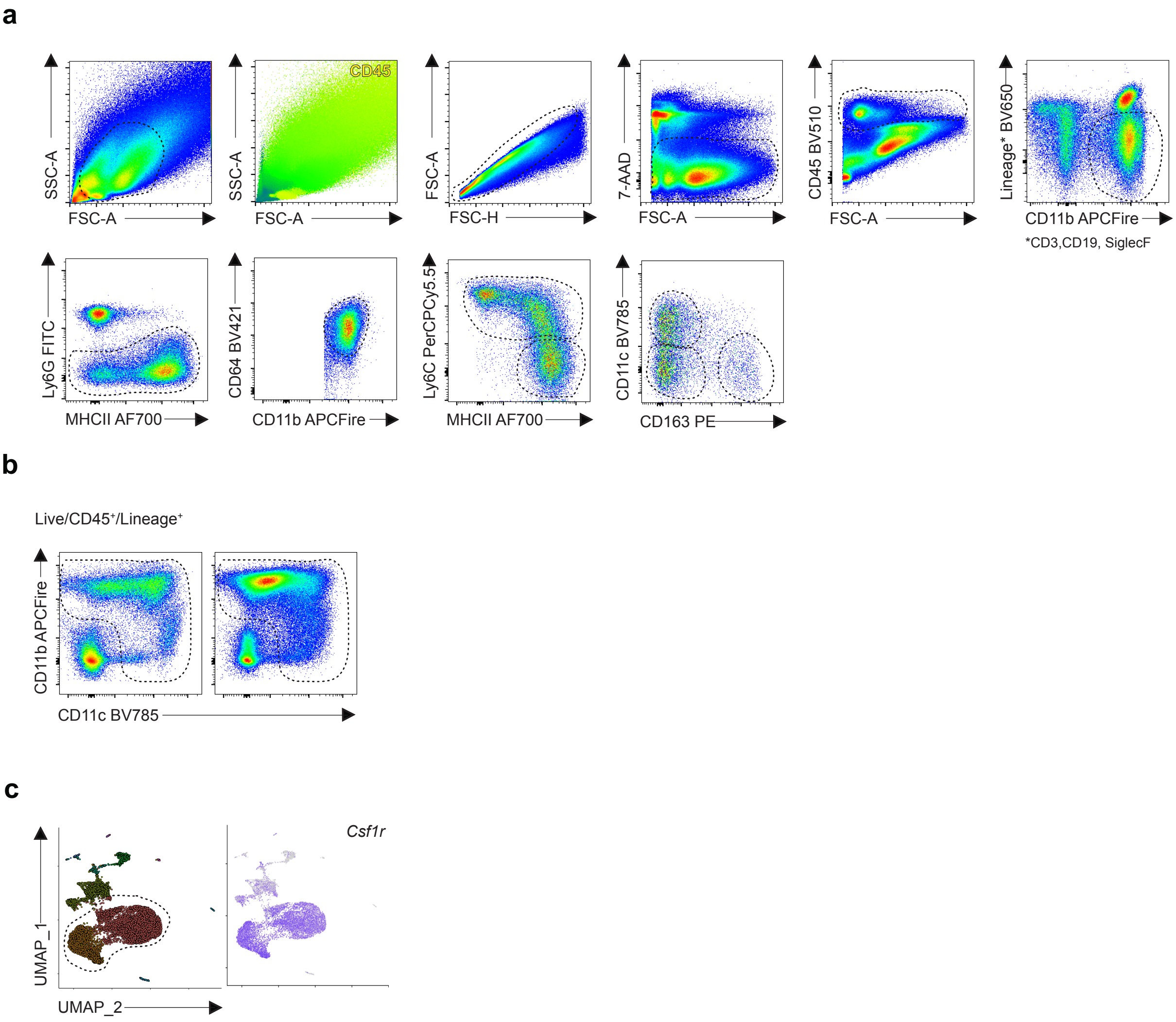

**Figure S2 Mouse flow cytometry macrophage identification strategy.**

**a)** Representative flow cytometry analysis of whole colon digest, obtained from day6 2% DSS treated C57BL/6J (WT) mice.

**b)** Flow cytometry purified (FACS) CD11b/c$^{+}$ cells for scRNA-seq from steady state (left) or day6 2% DSS (right) WT mice

**c)** UMAP dimensionality reduction analysis (left) using scRNA-seq from cells in b), with *Csf1r*$^{+}$ cell selection (Featureplot of gene expression, (right)) used for downstream analysis.

### Figure S3

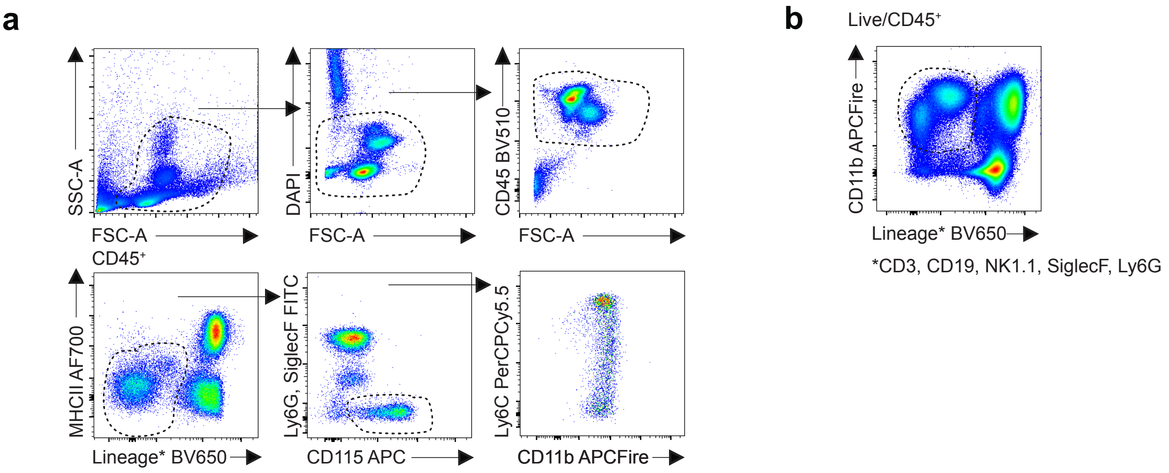

**Figure S3 Mouse flow cytometry Blood/BM identification strategy.**

**a)** Representative flow cytometry analysis of whole blood, obtained from 8-12week old C57BL/6J (WT) mice.

**b)** Flow cytometry purified (FACS) CD11b$^{+}$ cells for scRNA-seq from steady state or day6 2% DSS WT mice

**c)** Representative flow cytometry analysis and tdTomato expression of bone marrow, obtained from tibia/femur of 8-12week old *Ms4a3*$^{Cre/+}$.*Rosa26*$^{CAG\text{-}LSL\text{-}tdTomato/+}$ mice.

#### Figure S4

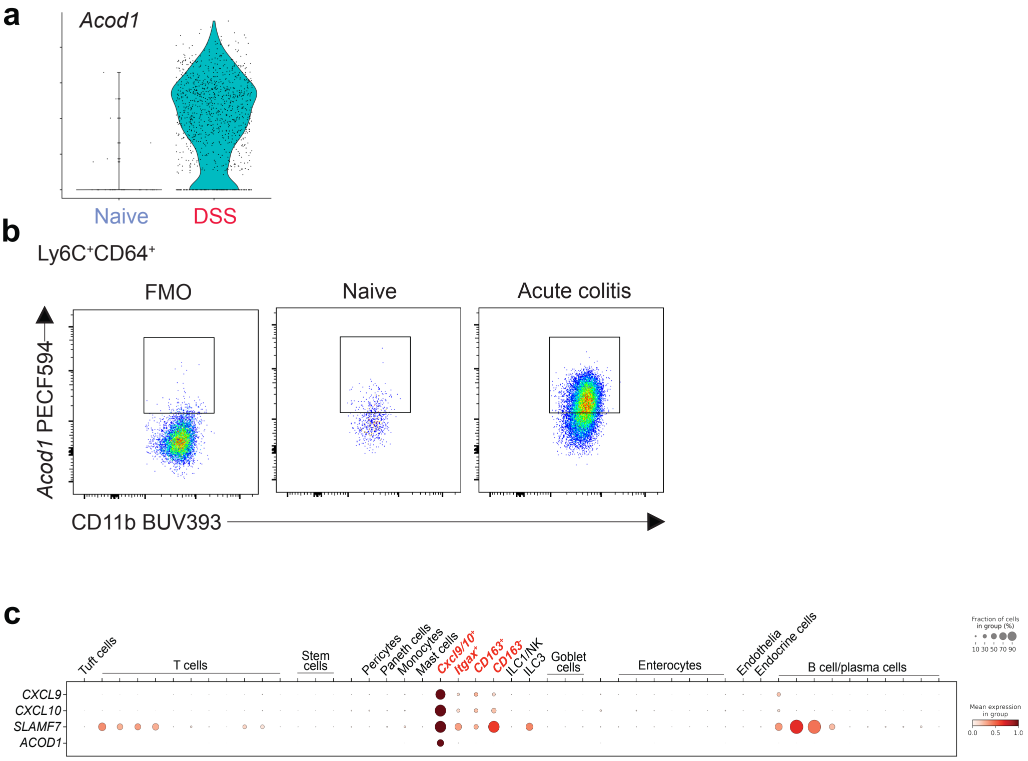

**Figure S4 Mouse flow cytometry Blood/BM identification strategy.**

**a)** Violin plot of *Acod1* expression in *Ccr2*$^{+}$ cells derived from scRNA-seq of colon CD11b/c$^{+}$ cells from steady state or day 6 2% DSS treated 8-12week old C57BL/6J (WT) mice from Figure 2a)

**b)** Representative flow cytometry analysis of *Acod1* probe mRNA (Primeflow) or fluorescence minus one control (FMO) in CD64$^{+}$Ly6C$^{+}$ monocytes obtained from steady state or day 6 2% DSS treated C57BL/6J (WT) mice

**c)** Bubble plot of selected genes across all atlased cell types from scRNA-seq analysis of 121 mucosal biopsies from endoscopically inflamed (active) ileal Crohn's disease and healthy controls.

### Figure S5

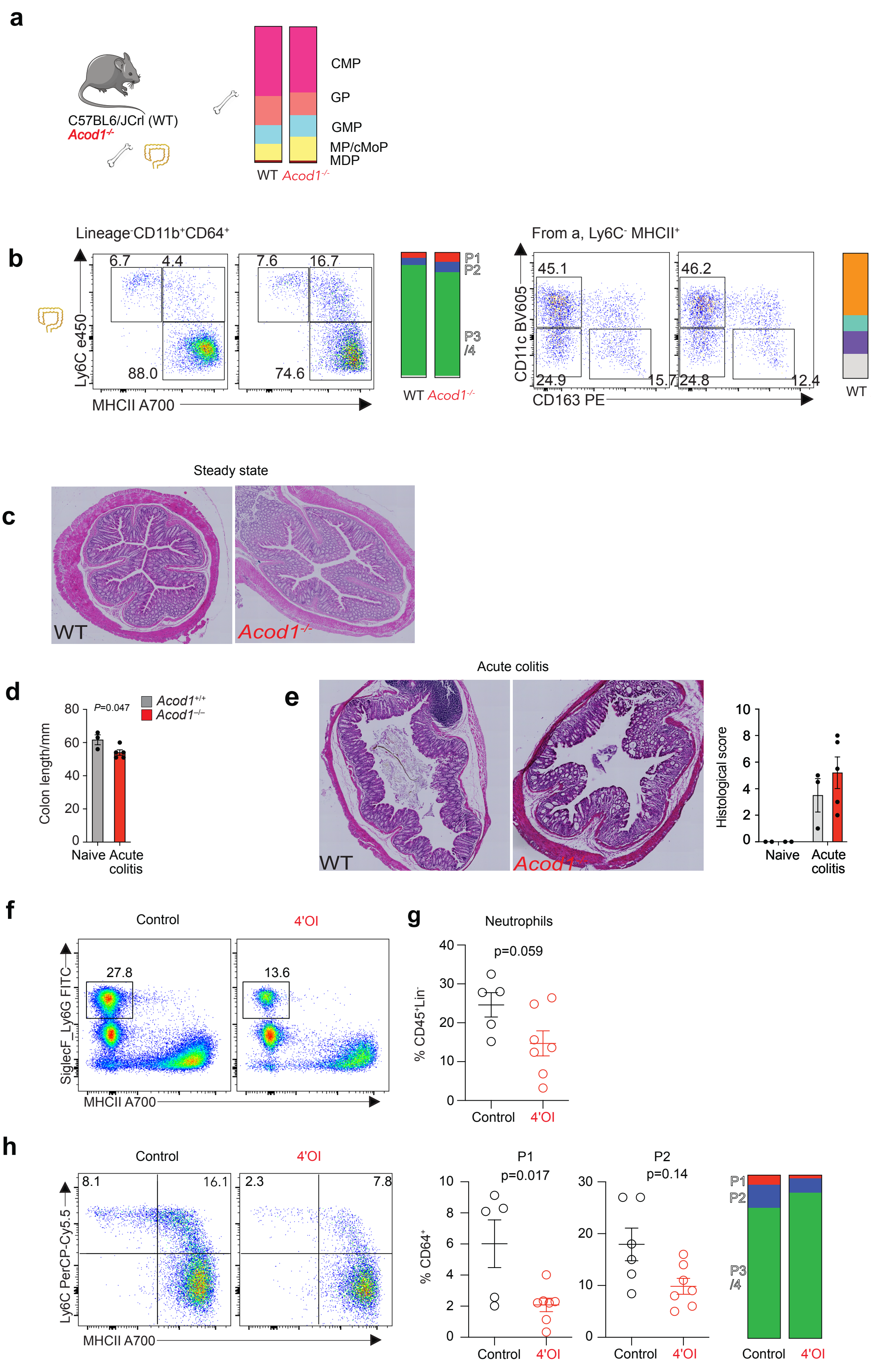
a
C57BL6/JCrl (WT)
Acod1-/-
CMP
GP
GMP
MP/cMoP
MDP
WT Acod1-/-
b
Lineage-CD11b+CD64+
Ly6C e450
MHCII A700
6.7
4.4
88.0
7.6
16.7
74.6
P1
P2
P3/4
WT Acod1-/-
From a, Ly6C- MHCII+
CD11c BV605
CD163 PE
45.1
24.9
15.7
46.2
24.8
12.4
CD11c
CD163
DN
Other
WT Acod1-/-
c
Steady state
WT
Acod1-/-
d
Colon length/mm
P=0.047
Acod1+/+
Acod1−/−
Naive
Acute colitis
e
Acute colitis
WT
Acod1-/-
Histological score
Naive
Acute colitis
f
Control
4'OI
27.8
13.6
SiglecF_Ly6G FITC
MHCII A700
g
Neutrophils
p=0.059
% CD45+Lin-
Control
4'OI
h
Control
4'OI
8.1
16.1
2.3
7.8
Ly6C PerCP-Cy5.5
MHCII A700
P1
p=0.017
P2
p=0.14
% CD64+
P1
P2
P3/4
Control
4'OI

**Figure S5 Endogenous itaconate (*Acod1*$^{-/-}$) deficiency in steady state and experimental colitis and exogenous synthetic itaconate supplementation to C57BL/6 (WT) mice**

**a)** Stack bar chart of CD117$^{+}$ bone marrow progenitors, using the flow cytometry identification strategy from Supp. Figure 3c and Yanez et al., comparing steady state 8-16week old *Acod1*$^{+/+}$ and *Acod1*$^{-/-}$ mice.

**b)** Representative flow cytometry analysis of CD64$^{+}$ monocytes/macrophages from steady state 8-16week old *Acod1*$^{+/+}$ and *Acod1*$^{-/-}$ mice. Stack bar chart of CD64$^{+}$ Ly6C$^{+}$MHC-II$^{-}$ (P1), Ly6C$^{+}$ MHC-II$^{+}$ (P2) and Ly6C$^{-}$ MHC-II$^{+}$ (P3/4)(left) and P3/4 cells, split by CD11c, CD163 or double negative (DN) expression (right).

**c)** Representative transverse sections of distal colon, H&E stained from steady state 8-16week old *Acod1*$^{+/+}$ and *Acod1*$^{-/-}$ mice.

**d)** Colon length from day6, 2% DSS treated, 8-16week old *Acod1*$^{+/+}$ and *Acod1*$^{-/-}$ mice. Data from 1 experiment, representative from 3 independent experiments and 18mice. Data analysed by unpaired Student's *t* test.

**e)** Representative transverse sections of distal colon, H&E stained from either steady state or day6, 2% DSS treated 8-16week old *Acod1*$^{+/+}$ and *Acod1*$^{-/-}$ mice(left) with histological score (Ho et al. 2018). Data from 1 experiment, representative from 3 independent experiments and 18mice.

Representative flow cytometry analysis of whole colon digest from C57BL/6J (WT) mice treated with 4 days 2% DSS, then 14 days normal drinking water, comparing 48hour administration of intraperitoneal 50mg/kg 4-octyl-itaconate or vehicle control. **f)** and **g)** Representative CD11b$^{+}$Ly6G$^{+}$ neutrophil flow cytometry plots and summary data (proportion of parent CD11b$^{+}$Lineage$^{-}$ cells). **h**) Representative CD64$^{+}$ monocyte/macrophage flow cytometry plots, summary data and stacked bar chart of CD64$^{+}$ cells (P1,P2,P3/4 proportion of parent CD64$^{+}$ cells). Data from 1 experiment, representative from 2 independent experiments and 28mice. Data analysed by unpaired Student's *t* test.

| | Healthy | CD |
|---|---|---|
| N | 71 | 50 |
| Age (mean±SD) | 55(15) | 39(13) |
| Male (%) | 31(44) | 24(48) |
| On IBD Medication (%) | 0 | 25(50) |
| Anti-TNF | NA | 8(16) |
| Immunosupressants (IM) | NA | 7(14) |
| Anti-TNF + IM | NA | 5(10) |
| Corticosteroids | NA | 3(6) |
| Ustekinumab | NA | 1(2) |
| 5'ASA | NA | 1(2) |

**Supplementary Table 1; Participant characteristics for scRNA-seq**

| | Healthy | IBD |
|---|---|---|
| N | 11 | 30 |
| Age (median±IQR) | 50(39) | 42(25) |
| Female (%) | 3 (27) | 13(43) |
| IBD subtype | | |
| Ulcerative colitis (n/%) | NA | 11 (37) |
| Crohn's disease (n/%) | NA | 13 (43) |
| IBD-unclassifed (n/%) | NA | 6 (20) |
| Endscopic activity (biopsies) | | |
| Uninflamed (n) | NA | 38 |
| Inflamed (n) | NA | 27 |
| Current smoker (n/%) | 0 | 4 (13) |
| Disease duration / yrs (median±IQR) | NA | 12(19) |
| CRP (median±IQR) | 1(4) | 9(17) |
| Albumin (median±IQR) | 41(4) | 36(6) |
| Haemoglobin (median±IQR) | 138(18) | 136(26) |
| Bio-naïve (n/%) | NA | 14(47) |

**Supplementary Table 2; Participant characteristics for flow cytometry**

| Flow cytometry | Clone | Supplier | Cat number |
|---|---|---|---|
| **Mouse** | | | |
| Rat monoclonal CD11b APC-Fire750 | M1/70 | Biolegend | 101262 |
| Armenian hamster monoclonal CD11c BV785 | N418 | Biolegend | 117336 |
| Rat CD16/32 TruStain FcX | 93 | Biolegend | 101320 |
| Rat monoclonal CD19 Biotin | 6D5 | Biolegend | 115504 |
| Rat monoclonal CD19 PE-Cy5 | 1D3 | Invitrogen | 15-0192-82 |
| Rat monoclonal CD3 Biotin | 17A2 | Biolegend | 100244 |
| Rat monoclonal CD3 PE-Cy5 | 17A2 | Biolegend | 100274 |
| Rat monoclonal CD45 BV510 | 30-F11 | Biolegend | 103138 |
| Mouse monoclonal CD45.1 FITC | A20 | Biolegend | 110706 |
| Mouse monoclonal CD45.2 AF700 | 104 | Biolegend | 109822 |
| Mouse monoclonal CD64 BV421 | X54-5/7.1 | Biolegend | 139309 |
| Rat monoclonal Ly6C PerCP-Cy5.5 | HK1.4 | Biolegend | 128012 |
| Rat monoclonal Ly6G Biotin | 1A8 | Biolegend | 127604 |
| Rat monoclonal Ly6G FITC | 1A8 | Biolegend | 127606 |
| Rat monoclonal MHCII (IA-IE) AF700 | M5/114.15.2 | Biolegend | 107622 |
| Rat monoclonal CD163 PE | S15049F | Biolegend | 156704 |
| Mouse monoclonal pro-IL-1β PerCP eFluor710 | NJTEN3 | Invitrogen | 46-7114-82 |
| Rat monoclonal CD319 APC | 4G2 | Biolegend | 152004 |
| Rat monoclonal CD274 PE | MIH7 | Biolegend | 155404 |
| Rat monoclonal CCRL2 BV605 | BZ2E3 | BDBiosciences | 743685 |
| Rat monoclonal TNF BV711 | MP6-XT22 | Biolegend | 506349 |
| Rat monoclonal CD34 BV421 | SA376A4 | Biolegend | 152208 |
| Rat monoclonal CD115 PECy7 | AFS98 | Biolegend | 135524 |
| Rat monoclonal CD135 APC | A2F10 | Biolegend | 135310 |
| Rat monoclonal CD16/32 APC/Cy7 | 93 | Biolegend | 101328 |
| Mouse monoclonal NK1.1 Biotin | PK136 | Biolegend | 108704 |
| Rat monoclonal Irg1 AF647 | *Primeflow* | Thermoscientific | VB1-11673-PF |
| **Human** | | | |
| Mouse monoclonal CD123 BB700 | 7G3 | BDBiosciences | 566482 |
| Mouse monoclonal CD38 BUV737 | HB7 | BDBiosciences | 612825 |
| Mouse monoclonal CD45 BUV395 | HI30 | BDBiosciences | 563792 |
| Mouse monoclonal CD1c BV786 | F10/21A3 | BDBiosciences | 742750 |
| Mouse monoclonal CD141 BV711 | 1A4 | BDBiosciences | 563155 |
| Mouse monoclonal CD163 BV650 | GHI/61 | BDBiosciences | 563888 |
| Mouse monoclonal CD14 BV421 | MφP-9 | BDBiosciences | 563743 |
| Mouse monoclonal CD55 PECy7 | JS11 | Biolegend | 311314 |
| Mouse monoclonal CD3 PE-CF594 | UCHT1 | BDBiosciences | 562280 |
| Mouse monoclonal CD19 PE-CF594 | HIB19 | BDBiosciences | 562294 |
| Mouse monoclonal CD11a APC/Fire750 | TS2/4 | Biolegend | 350612 |
| Mouse monoclonal HLA-DR AF700 | G46-6 | BDBiosciences | 560743 |
| Mouse monoclonal CD274 BV605 | 29E.2A3 | Biolegend | 329724 |
| Mouse monoclonal CD319 APC | 162.1 | Biolegend | 331810 |
| Mouse monoclonal CCRL2 PE | K097F7 | Biolegend | 358304 |
| Mouse monoclonal IL-1β FITC | JK1B-1 | Biolegend | 508206 |
| Mouse monoclonal TNF BV650 | MAb11 | Biolegend | 502938 |
| ***Other flow cytometry reagents*** | | | |
| 7-AAD | | Biolegend | 420404 |
| DAPI (4',6-diamidino-2-phenylindole, dihydrochloride) | | ThermoScientific | 62247 |
| Streptavidin BV650 | | Biolegend | 405232 |
| Zombie NIR Fiable Viability Dye | | Biolegend | 423106 |
| SYTOX Green nucleic acid stain | | ThermoScientific | S7020 |

| | | | |
|---|---|---|---|
| **Immunofluroescent imaging** | | | |
| Rabbit monoclonal ACOD1 (IRG1) | | Cell Signalling Technology | 17805 |
| Goat monoclonal IBA1 | | NOVUSBio | NB100-1028 |

**Supplementary Table 3; Antibody reagent details**

| Gene | Forward | Reverse |
|---|---|---|
| *Actb* | GGCTGTATTCCCCTCCATCG | CCAGTTGGTAACAATGCCATGT |
| *Acod1* | GCTTTTGTTAATGGTGTTGCTG | GGCTTCCGATAGAGCTGTGA |
| *Cxcl9* | TCGGACTTCACTCCAACACAG | AGGGTTCCTCGAACTCCACA |
| *Cxcl10* | TCTGAGTGGGACTCAAGGGAT | AGGCTCGCAGGGATGATTTC |
| *Tnf* | CCCTCACACTCAGATCATCTTCT | GCTACGACGTGGGCTACAG |
| *Il1b* | TGGACCTTCCAGGATGAGGACA | GTTCATCTCGGAGCCTGTAGTG |

**Supplementary Table 4; Mouse RT-PCR primer details**
